# Discovering and exploring the hidden diversity of human gut viruses using highly enriched virome samples

**DOI:** 10.1101/2024.02.19.580813

**Authors:** Moreno Zolfo, Andrea Silverj, Aitor Blanco-Míguez, Paolo Manghi, Omar Rota-Stabelli, Vitor Heidrich, Jordan Jensen, Sagun Maharjan, Eric Franzosa, Cristina Menni, Alessia Visconti, Federica Pinto, Matteo Ciciani, Curtis Huttenhower, Anna Cereseto, Francesco Asnicar, Hiroaki Kitano, Takuji Yamada, Nicola Segata

## Abstract

Viruses are an abundant and crucial component of the human microbiome, but accurately discovering them via metagenomics is still challenging. Currently, the available viral reference genomes poorly represent the diversity in microbiome samples, and expanding such a set of viral references is difficult. As a result, many viruses are still undetectable through metagenomics even when considering the power of *de novo* metagenomic assembly and binning, as viruses lack universal markers. Here, we describe a novel approach to catalog new viral members of the human gut microbiome and show how the resulting resource improves metagenomic analyses. We retrieved >3,000 viral-like particles (VLP) enriched metagenomic samples (viromes), evaluated the efficiency of the enrichment in each sample to leverage the viromes of highest purity, and applied multiple analysis steps involving assembly and comparison with hundreds of thousands of metagenome-assembled genomes to discover new viral genomes. We reported over 162,000 viral sequences passing quality control from thousands of gut metagenomes and viromes. The great majority of the retrieved viral sequences (∼94.4%) were of unknown origin, most had a CRISPR spacer matching host bacteria, and four of them could be detected in >50% of a set of 18,756 gut metagenomes we surveyed. We included the obtained collection of sequences in a new MetaPhlAn 4.1 release, which can quantify reads within a metagenome matching the known and newly uncovered viral diversity. Additionally, we released the viral database for further virome and metagenomic studies of the human microbiome.

## Introduction

Viruses are the most abundant biological entity on Earth and are key players in many environments ^1,2^, including the human gut microbiome ^3^. Bacteriophages (or simply “phages”), in particular, make up the majority of the human gut viruses, the so-called human gut “virome” ^4,5^. Phages have been associated with shifts in the resident microbial community in the gut, both as facilitators of lateral gene transfer ^6^ and as shapers of microbial communities ^7,8^. Phages have been implicated in many conditions and diseases including inflammatory bowel disease ^9,10^, malnutrition ^11^, cancer ^12^, and diabetes ^13^, but may also have a crucial role in promoting and maintaining gut and systemic health ^14^. Viromes also have the potential to be targeted modulators of the gut ecosystem, although their use as reproducible therapeutic tools is still far from being possible on a large scale ^15^.

Metagenomics ^16–18^ enabled the study of the human microbiome without the need for cultivation through shotgun next-generation sequencing (NGS). Since experimental isolation of viruses represents a formidable challenge, the use of NGS approaches is key to fully catalog the existent viral diversity. However, while metagenomics can provide a census of the human microbiome at an unprecedented depth, the metagenomic identification of viruses is still daunting. The biggest limitations in sequence-based viral characterization are the huge diversity of viruses and the lack of universal viral genomic markers, which hinder the identification of viral entities from raw sequencing reads. Numerous computational tools were developed to enable viral detection from metagenomes ^19–24^, some of which are also capable of *de novo* viral prediction. However, all these approaches rely, to some extent, on the information available from previously characterized viruses. This is true both for tools that search for direct matches against viral genes or proteins ^19,23,25,26^, as well as for those that exploit viral genetic patterns or use trained machine learning models to predict viruses ^27,28^.

Developing high-quality genomic catalogs of viral genomes is therefore crucial as a fundamental step in phage description and as a reliable and comprehensive reference for NGS virome studies. To date, 18,719 viral genomes are available in public repositories such as RefSeq ^29^. Most of them belong to model viruses or to pathogens of established clinical relevance. While several studies exploited sequence assembly to reconstruct genomes (metagenome-assembled genomes, or MAGs) from metagenomes, and released atlases of unprecedentedly large bacterial and archaeal diversity ^30–33^, these approaches rely on steps such as contig binning ^34,35^ and marker-based quality control ^36^ that are less exploitable on viral genomes. Such computational limitations for the *de novo* identification of divergent viral sequences (the so-called “viral dark matter”) can be partially overcome by experimental viral-like particle (VLP) enrichment protocols ^4,37,38^. VLP enrichment combines filtrations, concentrations, and selective gradient-purification to produce a sample composed mainly of viral sequences (i.e., a virome), but viral enrichment can have widely different efficiencies, even within-batch ^39^. While poorly enriched viromes can be useful for known virus detection and profiling, highly enriched viromes are crucial for *de novo* viral discovery. Overall, new tools and approaches are needed to expand the current knowledge of the virome fraction of human- and non-human-associated microbial systems.

In this study, we exploited highly enriched viromes to drive the *de novo* discovery of potentially viral sequences from metagenomes and viromes. We considered more than 3,000 viromes and selected those with high-purity to reconstruct high-confidence reference-free phages that underwent a series of screening and quality-control steps involving many thousands of metagenomes, low-enrichment viromes, known prokaryotic genomes, and collections of metagenome-assembled genomes. We obtained a catalog of 162,876 sequences that were clustered into 3,944 Viral Sequence Clusters (VSCs), most of which (85.1% of the total) represented potentially uncharacterized new viruses. We then surveyed the genetic and phylogenetic diversity, prevalence, and host-specificity (via CRISPR-Cas9 spacers’ matching) of such viral entities across many thousands of gut microbiome samples and showed that the generated catalog can serve as a computational base for viral detection. The catalog has been integrated into a new release of MetaPhlAn 4.1 ^40^ and both are available at https://segatalab.cibio.unitn.it/tools/metaphlan/.

## Results

To retrieve novel viral genomes and expand current phage databases for the gut microbiome, we developed a novel strategy exploiting the increased availability of both metagenomically sequenced VLP-enriched samples (viromes) and unenriched metagenomic samples (**Fig. 1A**). First, we retrieved 3,044 publicly available viromes from 49 datasets available in NCBI-SRA (see **Methods**, **Supplementary Table 1**). Under the assumption that a higher viral enrichment correlates with a lower quantitative detection of bacterial, archaeal, and eukaryotic taxa, we used ViromeQC ^39^ to estimate the viral enrichment level of each sample. We then considered contigs assembled from those viromes with the highest enrichment efficiency as highly trusted viral candidates (highly enriched viral contigs, HEVCs). We then extended HEVCs by screening more than 10,000 metagenomic samples and the associated set of >255M contigs and >150,000 *de novo* reconstructed bacterial taxa.

**Figure 1.**
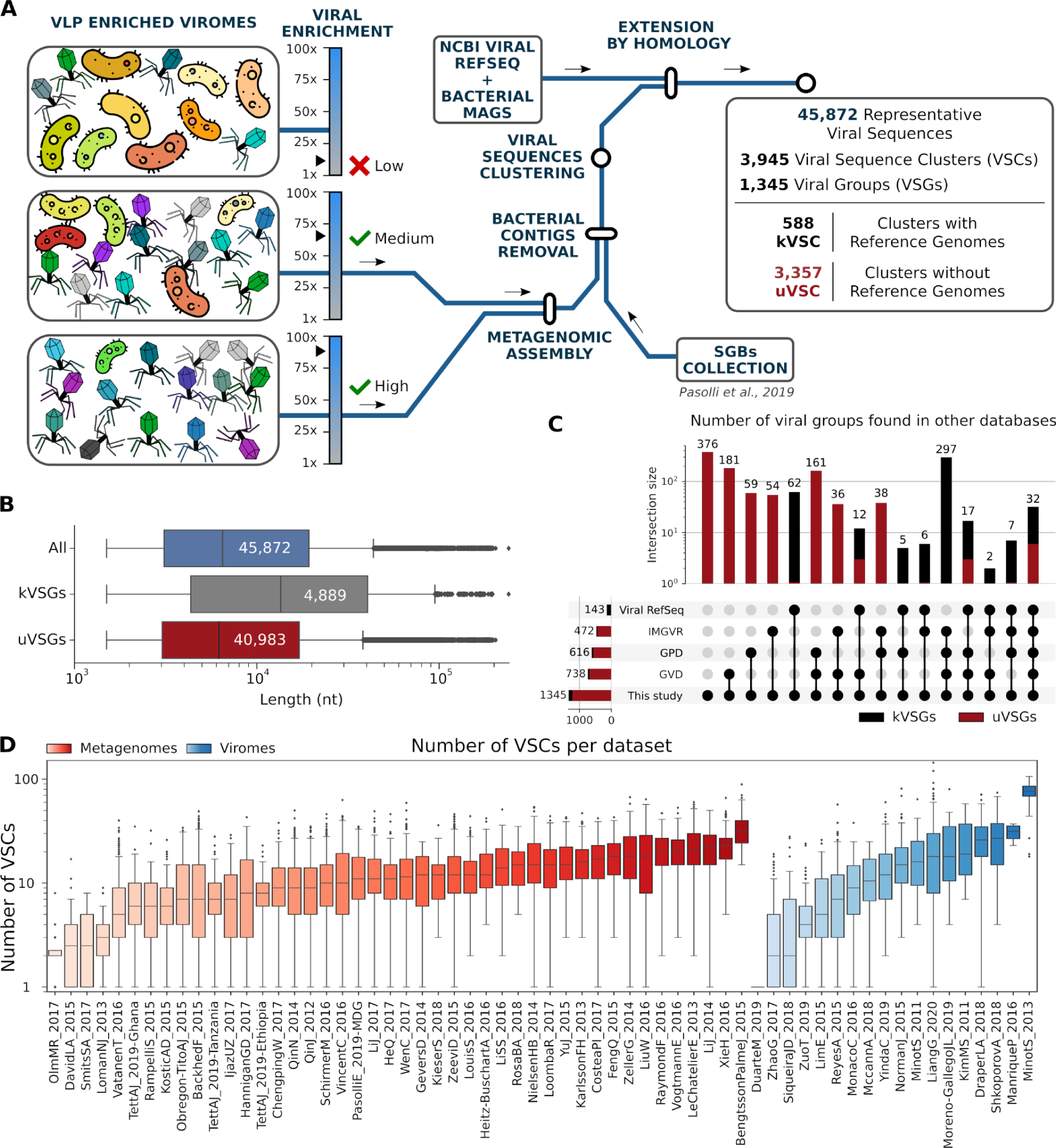
Methodology and resource overview. **(A)** Contigs were assembled from 255 highly enriched viromes (ViromeQC VLP enrichment >50x, see **Methods** and **Supplementary Table 1**). The resulting 120,041 contigs were screened to remove residual non-viral contaminants, and clustered at 90% identity. Clusters were then expanded with close sequences from 9,428 metagenomes and 2,789 lower-enrichment viromes. A set of 45,872 sequences (3,944 viral clusters) were selected as representatives.Viral Sequence Groups (VSGs) were formed via a graph clustering approach based on sequence similarity. **(B) Lengths of the viral representative sequences.** Distribution of the sequence length of the 45,872 viral representative sequences (blue box), stratified by sequences belonging to known (gray) and unknown (red) Viral Sequence Groups (VSGs). VSGs are groups of similar Viral Sequence Clusters (VSCs) clustered together by a graph-based approach. Groups were labeled as known when they contained at least one reference genome from Viral RefSeq, and unknown otherwise. **(C) Number of Viral Sequence Groups found in other viral databases.** The upset plot represents the intersection sizes (i.e., number of VSGs) found in any combination of four viral catalogs (RefSeq, IMG/VR, GPD, and GVD). The red bar indicates unknown VSGs. **(D) Number of viral sequences found in assemblies of stool samples, grouped by dataset.** Boxes encompass distribution quartiles, whiskers extend to 1.5 IQR. Red-shaded boxes are gut metagenomes, while blue-shaded boxes indicate VLP-enriched viromes. Datasets are described in **Table 1**. Datasets with samples from multiple sources (e.g., stool + oral metagenomes) are not included in the plot.

**Table 1.** Assembled contigs from the 255 highly-enriched viromes. Samples with a ViromeQC enrichment score >50x were considered to contain predominantly viral sequences. Samples are grouped by dataset and sample type. HQ reads refer to the number of reads that were retained after the quality-control step. Additional data for each sample is available in **Supplementary Table 1**.

| Dataset (Reference) | Sample Type | Samples | HQ Reads | Contigs | Median VQC score | Median N50 | Median Contig Length | Max Contig Length |
| --- | --- | --- | --- | --- | --- | --- | --- | --- |
| LiangG_2020 <sup>43</sup> | Human Stool | 71 | 4.82E+08 | 59154 | 100 | 5266 | 890 | 175037 |
| ReyesA_2015 <sup>11</sup> | Human Stool | 45 | 2.34E+06 | 10019 | 87 | 2589 | 893 | 44781 |
| LyM_2016 <sup>44</sup> | Human Stool | 43 | 2.23E+07 | 12345 | 100 | 3009 | 875 | 98074 |
|  | Human Oral | 21 | 9.84E+06 | 6201 | 100 | 3729 | 1.109 | 38713 |
| LimE_2015_MDA <sup>45</sup> | Human Stool | 17 | 2.21E+06 | 410 | 100 | 5440 | 969 | 84665 |
| MinotS_2013 <sup>46</sup> | Human Stool | 12 | 3.65E+08 | 27148 | 100 | 9221 | 789 | 393171 |
| NormanJ_2015 <sup>9</sup> | Human Stool | 6 | 1.51E+07 | 3195 | 63.5 | 2068 | 733 | 124399 |
| DeCarceraDA_2015 <sup>47</sup> | Freshwater | 5 | 5.30E+07 | 46462 | 95 | 1538 | 955 | 85920 |
| ArkipovaK_2017 <sup>48</sup> | Freshwater | 4 | 2.05E+07 | 281233 | 70.5 | 1061 | 654 | 140124 |
| PannarajP_2018 <sup>49</sup> | Human Stool | 4 | 3.86E+07 | 2196 | 100 | 6722 | 948 | 71301 |
| RouxS_2016 <sup>50,51</sup> | Ocean | 4 | 3.24E+08 | 706498 | 78.5 | 1864 | 835 | 200689 |
| LimE_2015_SIA <sup>45</sup> | Human Stool | 3 | 5.43E+05 | 120 | 100 | 1129 | 829 | 6039 |
| Moreno-GallegoJL_2019 <sup>52</sup> | Human Stool | 3 | 7.07E+06 | 3351 | 72 | 11143 | 702 | 187123 |
| PannarajP_2018 <sup>49</sup> | Human Milk | 3 | 2.34E+07 | 1785 | 96 | 4761 | 1.025 | 71301 |
| RosarioK_2009 <sup>53</sup> | Reclaimed Water | 3 | 7.30E+05 | 12407 | 100 | 967 | 720 | 19416 |
| PintoF_2020 (unpublished) | Freshwater | 2 | 8.97E+07 | 103294 | 61.5 | 2204 | 874 | 80726 |
| DeCarceraDA_2016 <sup>54</sup> | Freshwater | 2 | 2.57E+05 | 2579 | 100 | 1040 | 787 | 4264 |
| KimMS_2011 <sup>55</sup> | Human Stool | 2 | 2.20E+05 | 1235 | 92.5 | 2138 | 847 | 24667 |
| MinotS_2011 <sup>56</sup> | Human Stool | 2 | 8.47E+04 | 868 | 57 | 2501 | 821 | 31906 |
| RouxS_2012 <sup>57</sup> | Freshwater | 2 | 1.25E+06 | 46685 | 79.5 | 1062 | 761 | 27163 |
| LopezBueno_2009 <sup>58</sup> | Freshwater | 1 | 6.13E+05 | 1817 | 76 | 1075 | 773 | 6622 |
| **Total** |  | **255** | **1.5E+09** | **1.3E+06** |  |  |  |  |

### Selection of contigs from highly enriched human gut viromes

After uniform preprocessing to remove low-quality and short reads (see **Methods**), viromes were screened with ViromeQC ^39^ to determine their viral enrichment. Out of 3,044 viromes, the 255 samples with an enrichment score higher than 50x were considered “highly enriched” and were subjected to metagenomic assembly (see **Methods**). In total, 1.5 billion reads were assembled, producing 1.3M contigs longer than 500 nucleotides (**Table 1, Supplementary Table 1**), of which 120,041 originated from human gut viromes. Contigs were generally short (overall median length = 860, median N50 = 3,863, **Fig. 1B**), with 8,488 contigs longer than 5 kbp (7.1%) and 3,258 contigs longer than 10 kbp (2.7%).

We found that only 3,932 out of 120,041 contigs (3.28%) aligned to one viral genome in RefSeq ^41^ (>80% identity in 50% of the target, with aligned segments >= 500nt) (**Supplementary Table 2**). Moreover, 23,240 contigs (19.36%) had at least one hit against a collection of representative reference bacterial and archaeal genomes ^32,42^, and 34,424 (28.68%) aligned against at least one of the non-viral MAGs (i.e., prokaryotic MAGs of at least medium-quality) from Pasolli *et al*. ^32^. As viral sequences could match against bacterial genomes because of prophagic regions and MAGs could erroneously include non-bacterial contigs, it was intriguing to find that 81,212 (67.65%) contigs did not match against any sequence available as reference-based or reference-free cataloged microbial diversity. Moreover, several contigs were found multiple times in different viromes (**Supplementary Fig. 1**), with 7,607 contigs (6.34%) found in more than one dataset, and 1,801 contigs (1.5%) found in at least 4 datasets, supporting their biological validity and highlighting their potential prevalence in the human gut microbiome.

### Refinement of the viral contigs to remove further non-viral contamination

To address the hits against non-viral databases, we further exploited the compendium of MAGs assembled from 9,428 metagenomes and available as species-level genome bins (SGBs) from Pasolli *et al.* to screen the 120,041 sequences from highly enriched gut viromes. First, we excluded 29,237 (24.36%) contigs that were found in medium- to high-quality MAGs assigned to bacteria or archaea in more than 50 metagenomes. Then, we considered both the sequences from Pasolli *et al.* that were not binned into any MAG (not even in low-quality bins) as well as the organization of bacterial and archaeal MAGs and genomes in species-level genome bins (SGBs). Based on these two sets, we kept as viral contigs only those found in the unbinned fraction of at least 20 samples to ensure we considered viruses of sufficient prevalence and supported by enough single-sample assemblies. Overall, this procedure selected 4,952 contigs that were therefore a) of potential viral origin, b) rarely detected in bacterial genomes and MAGs, and c) highly prevalent in metagenomes and viromes. Finally, we also added 699 full genomes of known and prevalent human gut bacteriophages from RefSeq to the set of 4,952 contigs. These 699 full phage genomes from RefSeq were selected as those with a known bacterial host as per their description in NCBI and that were found in at least 20 metagenomes from the unbinned fraction of Pasolli *et al*.. This brought the total number of phage sequences to 5,651, forming the set of Highly Enriched Viral Contigs (HEVCs).

### The newly identified viral sequences comprise phages not detected by other approaches

We contextualized the 5,651 HEVCs against four available computational viral prediction tools. These methods were developed to determine if an assembled contig is a virus using direct or indirect concepts of similarity with known viruses ^19–21,23^. VirFinder classified as viral the largest number of sequences (n=3,175), while VirSorter in default mode detected 1,000 viral contigs. VirSorter in decontamination mode classified 695 RefSeq genomes (99.4%) as viral and was in general agreement with the other tools (ViralVerify: 692, VirFinder: 632, Virsorter: 582, Seeker: 142, **Supplementary Table 3**). Overall, only a small fraction of our sequences were classified as viral by all tools (from 153 to 186 out of 5,651 depending on the settings for VirSorter, **Fig. 2**). More importantly, 1,052 contigs (18.62%) in our collection could not be classified as viral by any tool, suggesting that our catalog may capture a substantial fraction of novel viral diversity.

**Figure 2.**
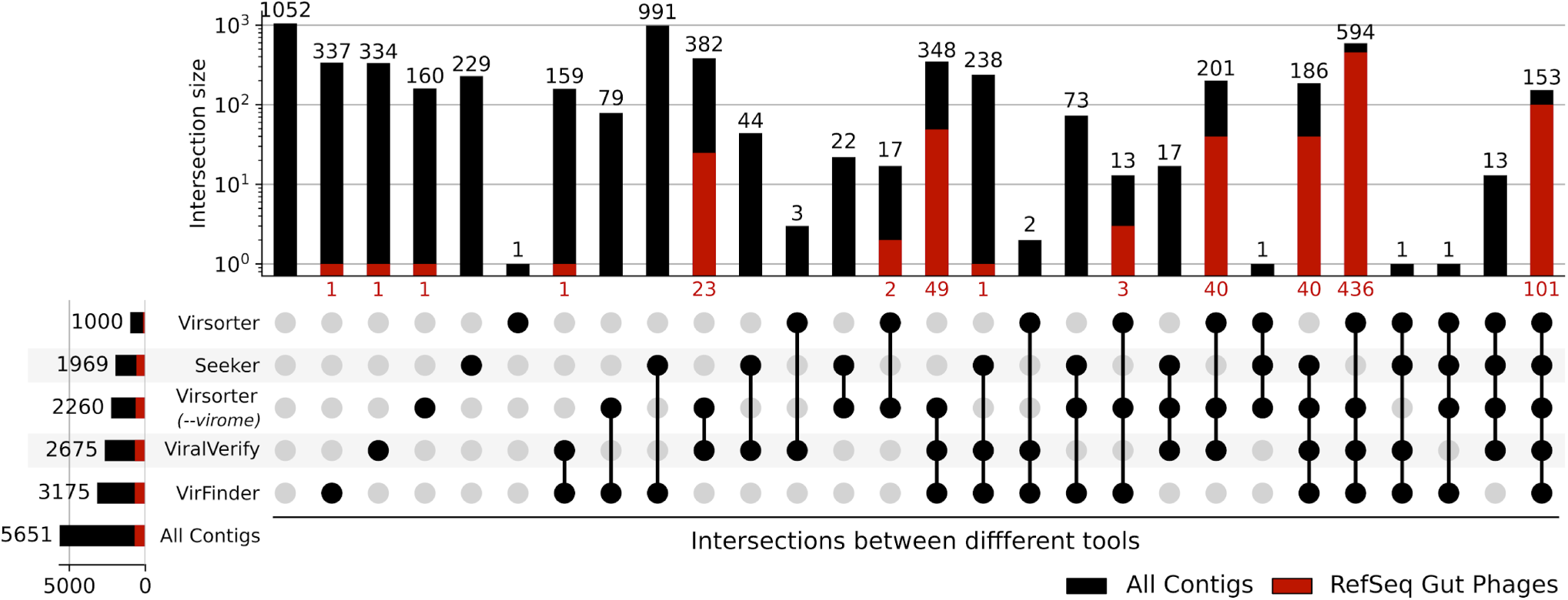
Comparison of different viral detection tools on the 5,651 contigs from RefSeq and highly-enriched gut viromes. The upset plot shows the number of contigs classified as viral by combinations of different tools (Virsorter, Seeker, ViralVerify, and VirFinder). The set of analyzed contigs includes 4,952 contigs from highly enriched viromes (black bars) and 699 reference genomes of human gut bacteriophages from RefSeq (red bars). The number of contigs in each intersection is shown on the top (overall number of contigs) and bottom (RefSeq reference gut bacteriophages) of each bar.

As an additional assessment, we applied the same four tools to a random set of 200 bacterial MAGs and 2,619 RefSeq-verified bacteriophage genomes. Bacterial MAGs were expectedly less likely to be labeled as viral, but 7.9% of the bacterial contigs were labeled as viral by at least one tool. Moreover, only 31.1% of the 2,619 reference bacteriophages in RefSeq were predicted as viral by all tools (**Supplementary Fig. 2**). This suggests that several phage sequences are still transparent to available viral classifiers because of their high novelty and divergence to any known virus.

Overall, these results highlight that our resource, which is the only one available that does not rely on already known phages, contains novel viral diversity, a substantial fraction of which is not identifiable by available methodologies.

### Building and expanding the viral sequence clusters (VSCs)

Because some of the retrieved viral contigs were highly similar, we clustered the 5,651 HEVCs at 90% identity with VSEARCH ^59^, obtaining 3,944 clusters that we call Viral Sequence Clusters (VSCs). Of these, 588 contained a viral reference genome and were labeled as known VSCs (kVSCs, 14.90%), and 3,357 were labeled as unknown (uVSC, 85.10%). On average, there were 1.38 sequences per cluster and the largest uVSC (c84, **Supplementary Table 3**) included 56 sequences. The largest kVSC (c1586) grouped 19 sequences, and included the reference genome of the PhiX174 Coliphage (NC_001422.1), underscoring the presence of undepleted NGS spike-in control in many metagenomes ^60^. The second and third largest clusters contained the sequences of widely prevalent gut bacteriophages crAssphage ^61^ (NC_024711), and fragments of *Bacteroides phage* B12414 (NC_0167701), respectively.

Unenriched metagenomes and viromes can still contain near-complete viral sequences. To access this additional diversity, we used Mash 2.0 ^62^ to search for sequences similar to those in the 3,944 VSCs against a) the contigs assembled from all the 3,044 viromes initially considered, and b) contigs in the unbinned fraction of Pasolli *et al.* (see **Methods**). This mapping performed at 90% sequence similarity threshold retrieved 157,225 additional contigs (126,894 from unbinned metagenomes, 30,331 from viromes, **Supplementary Table 3**) that were added to our proposed resource of 162,876 novel, high-confidence, and potentially viral sequences, recapitulated into the 3,944 VSCs. To enable easy profiling and re-use of the discovered viral sequences, while reducing unnecessary sequence redundancy, we selected 45,872 non-redundant representatives (see **Methods, Table 2**) for the VSCs. Sequences, metadata, and clustering data are publicly available for future studies (see **Data Availability**).

**Table 2.** Composition and origin of the 45,872 VSCs representative sequences. Summary of the 45,872 representatives of the total set of 162,876 sequences derived from viromes and metagenomes. Numbers in the brackets indicate how many sequences were sorted into unknown Viral Sequence Clusters (uVSCs). The selected cluster representatives are either the longest sequences within each 95% identity cluster, or sequences originating directly from highly-enriched viromes.

| Contig Source | Sequences | Avg Len. | St. Dev. Len. | Median Len. | Max Len. |
| --- | --- | --- | --- | --- | --- |
| Highly Enriched Viral Contigs (HEVCs) | 3,467 | 5,774.23 | 9,950.35 | 2,991 | 167,064 |
| of which in uVSCs | (3,418) | (5,620.47) | (9,263.16) | (2,978) | (167,064) |
| Viral RefSeq | 684 | 58,780.93 | 44,793.82 | 41,718 | 240,413 |
| Viromes | 8,804 | 12,853.21 | 20,580.28 | 5,615 | 204,229 |
| of which in uVSCs | (8,437) | (12,288.42) | (19,755.01) | (5,476) | (204,229) |
| Metagenomes | 32,917 | 20,923.09 | 30,367.33 | 7,592 | 198,476 |
| of which in uVSCs | (31,331) | (20,371.18) | (30,185.44) | (7,112) | (198,476) |
| **Any (Total)** | **45,872** | **18,793.83** | **28,758.97** | **6,485** | **240,413** |

### VSCs represent a large portion of previously unknown viral diversity

We evaluated the prevalence of each viral cluster within the datasets originally used to build the catalog. Out of the 3,944 VSCs, 339 (8.59%) were metagenomically assembled at least once in half of the gut metagenomic datasets (**Supplementary Table 4**). Among the most prevalent kVSCs, we detected the *crAssphage* (kVSC c72, detected in 45 datasets in at least one sample, average prevalence when detected 6.61%, s.d. 5.26%), as well as several *Enterobacteria* and *Shigella* phages (**Supplementary Fig. 3**). The fifth most prevalent viral cluster (kVSC c1586) was annotated as Coliphage phiX174, the spike-in control used on NGS platforms. This undepleted control sequence was found at least once in 15 out of 44 metagenomic datasets and in 10 out of 19 viral datasets. In the virome datasets where phiX174 was detected, it had an average prevalence of up to 60.4%, pointing at the pervasive nature of this undepleted spike-in control in public samples.

The majority of highly prevalent VSCs were uVSCs (i.e., no RefSeq viral genomes were present in the cluster). The three uVSCs with the highest average prevalence in gut datasets (uVSC c2353, c3062, c3861, **Supplementary Table 4**) attracted 8,838 sequences (8,333 of which from metagenomes, the others from viromes). Further investigation of these clusters indicated that sequences within each cluster were extremely similar, despite coming from different samples and datasets (median pairwise identity on multiple-sequence alignment >98.7%). None of the sequences matched against any artificial vector, linker, adapter, and primer in the NCBI sequence vector database (UniVec), and only 4.25% of the sequences in VSC c3062 matched against UniVec entries that could be both of natural and artificial origin (see **Methods**). Within these three VSCs (uVSCs c2353, c3062, c3861), the seven sequences from highly enriched viromes were not identified as viral by VirSorter, ViralVerify, Seeker, and VirFinder, nor they had matches against sequences in viral-RefSeq.

### CRISPR spacers analysis systematically reveals novel host-association of kVSCs and uVSCs

Clustered Regularly Interspaced Short Palindromic Repeats (CRISPR) spacers in bacterial genomes can be used to infer associations between viral hosts and microbes ^63^. The underlying assumption is that fragments of the bacteriophage genome can be integrated into the CRISPR locus when the target host is infected, and then be used as a proxy to determine if that phage is capable of infecting a given bacterial strain or species. To investigate the host associations of the viruses in our catalog, we predicted the CRISPR loci on a collection of 377,345 MAGs and 169,111 reference genomes, resulting in a total of 5,843,193 CRISPR spacers (see **Methods**). We then searched the CRISPR spacers in the VSCs and linked each viral cluster with the taxonomy of the corresponding MAGs. Most of the 3,944 VSCs (88.24%) had at least a match against a CRISPR spacer and the majority of them (74.7%) had a predicted host, while 494 VSCs (12.53%) had at least 5 associations with at least one microbial species, and 119 (3.02%) were associated with at least 10 different species, further validating the viral origin of VSCs (**Figure 3**, **Supplementary Table 5**). Moreover, 35 clusters were associated with more than 10 genera, and 17 of them with more than 3 phyla, spanning Actinobacteria, Firmicutes, and Bacteroidetes. The bacterial species most frequently associated with any of the VSCs were *Bacteroides uniformis*, *Parabacteroides distasonis*, and *Phocaeicola vulgatus*, which were associated with 220, 203, and 194 VSCs respectively.

**Figure 3.**
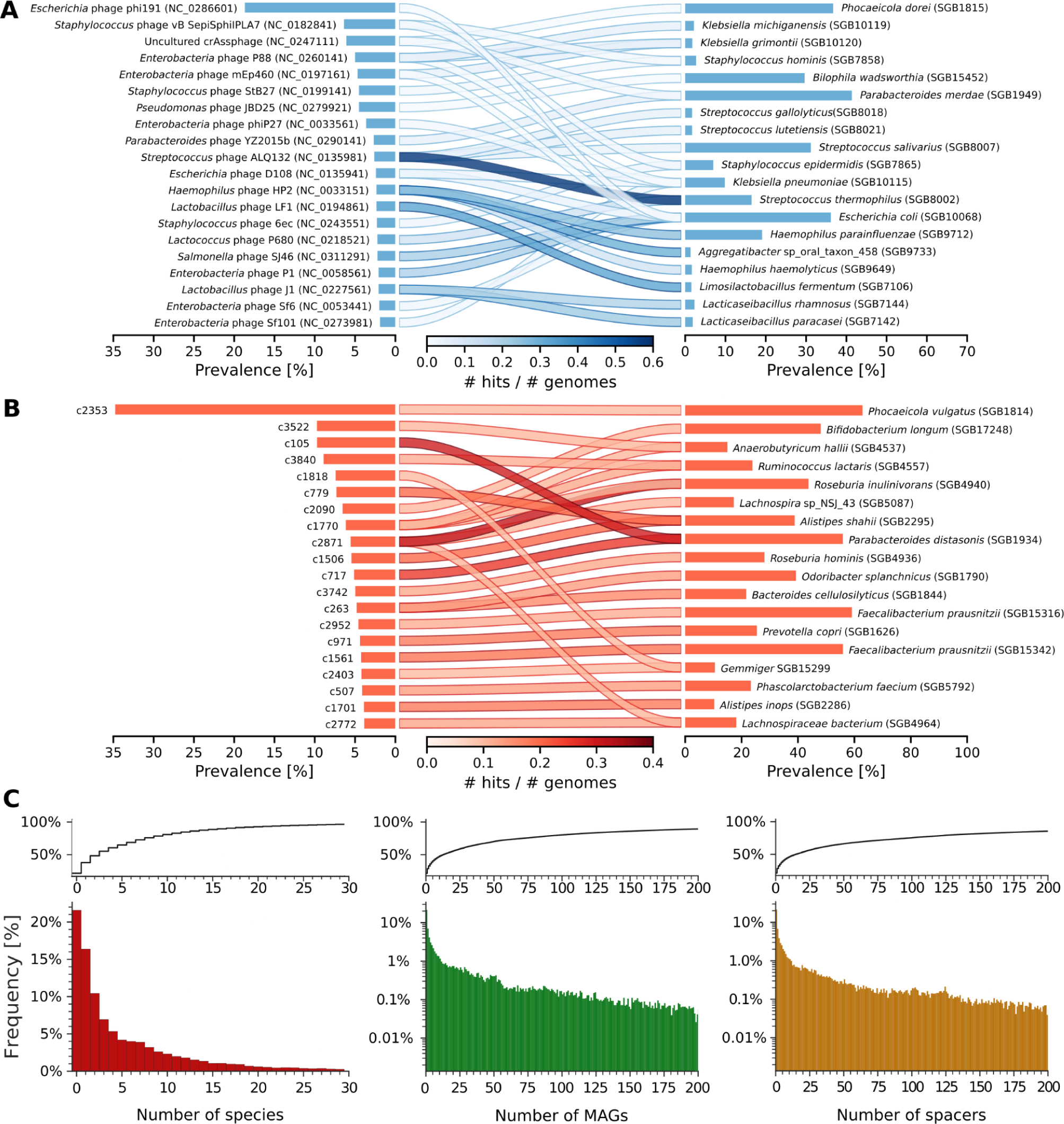
Association of bacteriophages clusters with microbial hosts via CRISPR spacers. The barplots on each side represent the prevalence of Viral Sequence Clusters (VSCs, left) and microbes (species-level genome bins, SGBs, right) from a set of 7,714 assembled metagenomes and 18,756 raw gut metagenomes. The top 20 known **(A)** and unknown **(B)** VSCs are shown. Viruses and hosts are connected if the viral cluster has >5 hits against a species confirmed in at least 0.5% of the genomes within the bin for known SGBs (**A**), or >10 hits against a species, confirmed in at least 10% of genomes in the bin for unknown SGBs (**B**). The shade of the connecting line represents the fraction of microbial genomes in the SGB for which a CRISPR-spacer hit is reported. Viral clusters without predicted hosts are not included, regardless of their prevalence. **(C) Association of Viral Groups and Hosts via CRISPR-spacers mapping.** Percentage of the 45,872 VSGs representative sequences (y-axis) that are associated with each number (x-axis) of distinct microbial species (red), MAGs (green), or CRISPR-spacers (orange). The cumulative distribution is reported above each histogram. CRISPR spacers were predicted from 377,346 Metagenomic Assembled Contigs and reference genomes (see **Methods**).

We compared our collection of viral sequences with other recently published databases of viruses and phages, with similar findings in terms of number of sequences and clusters being associated with bacterial hosts via CRISPR spacers. In particular, 91.8% of the 57,866 viral clusters of the Gut Phage Database ^64^ were associated with at least one species. Only 18.7% of the 935,363 vOTUs of IMG/VR (and 21.3% of the 2,377,994 individual sequences) ^65^ and only 27.2% of the 13,582 viruses in RefSeq were associated with a microbial species (**Supplementary Fig. 4**). This suggests that the VSCs collection contains as many, and in some cases more, sequences likely to be of phagic origin, while it has to be noted that this may also reflect that not all the databases are uniquely focused on bacteriophages (e.g., RefSeq or IMG/VR).

We then compared the prevalence of the VSCs with the prevalence of their putative host microbes in 18,756 gut raw metagenomes profiled with MetaPhlAn 4.1 (**see Methods**). Several highly prevalent viruses, both known and unknown, were associated with multiple prevalent and expected species of the gut microbiome. For example, kVSC c527 (containing the Streptococcus_phage_ALQ132 reference genome NC_013598.1) was associated with *Streptococcus thermophilus, S. lutetiensis, S. vestibularis, S. gallolyticus,* and *S. salivarius*. The kVSC c324 (a *Staphylococcus* phage based on available annotations) was associated with *S. hominis* and *S. epidermidis*, while several unknown and highly prevalent viruses were associated with microbes found in the human microbiome at prevalences >20% (**Figure 3**, **Supplementary Table 5**). uVSC c2353, the most prevalent unknown virus in the human gut assemblies, was associated with the highly prevalent *Phocaeicola vulgatus* (**Figure 3**), as well as *P. dorei, P. massiliensis*, and *P. coprocola*, but also similarly prevalent species of different genera such as *Parabacteroides distasonis*, *Bacteroides uniformis*, and *Bacteroides cellulosilyticus (***Supplementary Table 5**). Overall, this suggests a broad species tropism of the bacteriophages in the human gut, detectable when considering CRISPR-spacers associations. This is further corroborated by the detection of highly prevalent viruses that are associated in turn with likewise highly prevalent microbial species.

### Phylogenetic profiling of the sequence diversity within VSCs

Further analysis of the VSCs allowed us to reconstruct the phylogeny of several viruses. As an example, we analyzed known kVSC c718_c0 (containing *Lactococcus phage bIL67*, accession NC_001629, **Fig. 4A**) and kVSC c230_c0 (containing *Klebsiella phage KpnP*, accession NC_028670, **Fig. 4C**). We were also able to reconstruct the phylogeny of 262 sequences of kVSC c72_c0 (crAssphage cluster, **Supplementary Fig. 5**). The crAssphage-like sequences retrieved from highly-enriched viromes were all within the same clade, while contigs from viromes and metagenomes extended the phylogeny. This demonstrates that even unenriched samples can provide access to unseen viral diversity.

**Figure 4.**
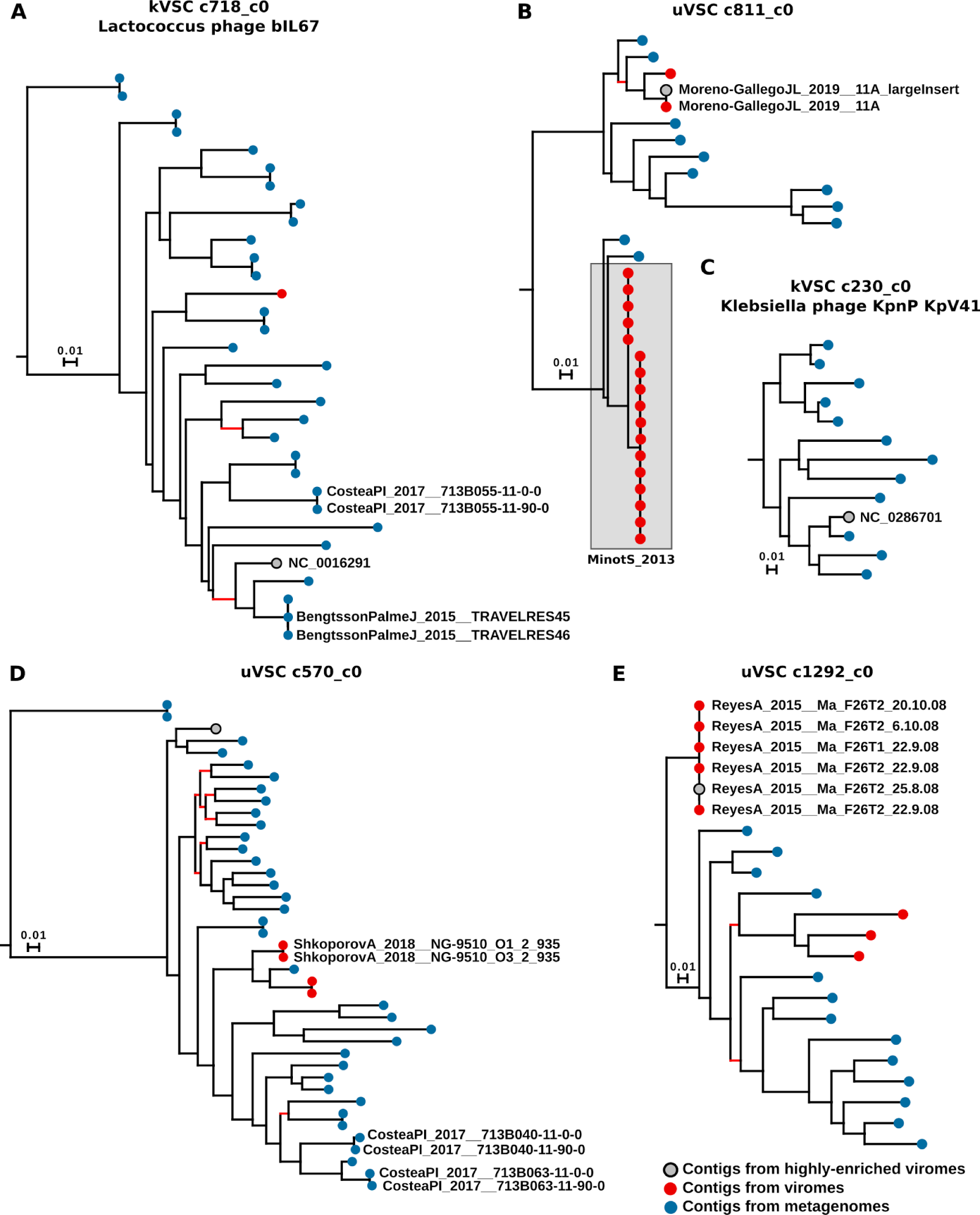
Phylogenetic trees of five prevalent Viral Sequences Clusters. A subset of the sequences in each cluster underwent multiple-sequence alignment and a Maximum Likelihood phylogenetic tree was generated. Gray nodes indicate contigs from the original collection of 5,651 contigs from highly enriched samples (HEVCs). Colored nodes indicate sequences selected by similarity to HEVCs from low-enrichment viromes (red nodes) and from unbinned metagenomes (blue nodes). Leaves labels indicate contigs retrieved from the same individuals (**A,B,D**), or from subjects sharing households (**E**). Branches with bootstrap confidence support lower than 75% are highlighted in red. Only sequences within 25% of the median sequence length of the whole cluster (**A,B,C**) or within 15% of the median length of the highly-enriched-sequences in the cluster (**D-E**) were aligned. Alignment lengths are provided in **Supplementary Table 6**.

When we incorporated the metadata of the original studies into the phylogenetic trees, we uncovered individual-specific phages that were also maintained in time. In the *Lactococcus* phage phylogeny, we identified at least two pairs of samples of the same individual taken at different time-points (**Fig 4A**). This was also seen in several other VSCs in longitudinal datasets ^43,46,66^ where multiple samples per individual were analyzed ^52,67^, or where individuals shared households ^11,43,44^. As an example, in the unknown cluster uVSC c811_c0, one clade contained sequences from the same sample sequenced with different library preparations (dataset Moreno-GallegoJL_2019), while the other clade was composed mainly of very similar sequences from the same individual sampled longitudinally (MinotS_2013, **Fig. 4B**). Also in uVSC c1292_c0, sequences from twins that shared the same household clustered in the same branch (ReyesA_2015, **Fig. 4E**). At least 38% of the Open Reading Frames of these three uVSCs matched a viral motif of V-FAM^68^ (see **Methods**), highlighting their very likely viral origin. Moreover, all uVSCs had partial matches with MAGs of gut-associated bacterial taxa as *Holdemanella biformis* (uVSC c811_c0) *Bifidobacterium adolescentis* (uVSC c570_c0), and *Prevotella copri* (uVSC c1292_c0). Both VirFinder and ViralVerify classified the three clusters as viral, while VirSorter classified only one (uVSC c1292) as a sure virus, and the other two as potential viruses (**Supplementary Table 6**).

The detection of the same sequence in samples of the same individuals highlights that not only some VSCs are prevalent across datasets, but also that they can persist and be shared within households (**Fig. 4E, Supplementary Fig. 6**). This is similar to what has been found by other studies, both for bacteria and viruses in the human microbiome ^52,69–72^. Finally, we also analyzed the phylogeny of the cluster that included the PhiX174 Coliphage. We noticed a complete absence of phylogenetic signals (median pairwise identity = 100%, average pairwise identity = 99.98%, n=1,133 sequences), confirming that such sequences were sequencing spike-in artifacts not filtered before metagenomic assembly.

### Integrating the new viral database with MetaPhlAn 4 reveals several highly prevalent unknown phages

To investigate the prevalence of viruses in publicly available human gut metagenomes, we included the 45,872 VSCs representatives into a new module of the MetaPhlAn 4.1 metagenomic taxonomic profiler ^40^. Viral sequences were grouped into 1,345 “Viral Sequence Groups” (VSGs, see **Methods**), to avoid closely related viruses to compete when attracting short sequencing reads. With this new module, the user can obtain the breadth and depth of coverage of each viral group, in addition to the default MetaPhlAn output. The viral profiling module for MetaPhlAn 4.1 is open source and available at http://segatalab.cibio.unitn.it/tools/metaphlan/ and as the bioconda package ‘metaphlan’, by using the option --profile_vsc.

We applied the newly developed module to detect viruses in 18,756 human gut metagenomes from 81 datasets, spanning healthy and diseased subjects across multiple geographic locations (**Supplementary Table 7**). We found several extremely prevalent clusters across multiple datasets, study cohorts, and countries (**Fig. 5**), with seven potential unknown viral groups found in more than 30% of the metagenomes. The most prevalent group was M697, a set of 29 contigs that had a combined prevalence of 74.87% (detected in 14,042 samples). M697 did not contain any RefSeq phage and was thus labeled as “unknown”, as well as the ten most prevalent viral groups. The most prevalent known viral groups were crAssphage (overall prevalence = 19.07%), *Enterobacteria Phage P2* (16.21%), and *Lactococcus Phage phi7* (12.97%). The viral group with the highest average per-dataset prevalence was the *Coliphage PhiX174*, highlighting once again the widespread presence of such contaminants in metagenomes, as it was detected in 25 out of 81 datasets. Generally, as expected, the prevalence of viral groups calculated by mapping raw reads to VSCs was higher if compared to mapping assembled contigs to VSCs. This indicates a higher sensitivity of the mapping-based approach implemented in MetaPhlAn for detecting VSCs (**Fig. 5**, **Supplementary Fig. 3**). Of note, the prevalence of crAssphage was comparable with the findings of other studies ^73^.

**Figure 5.**
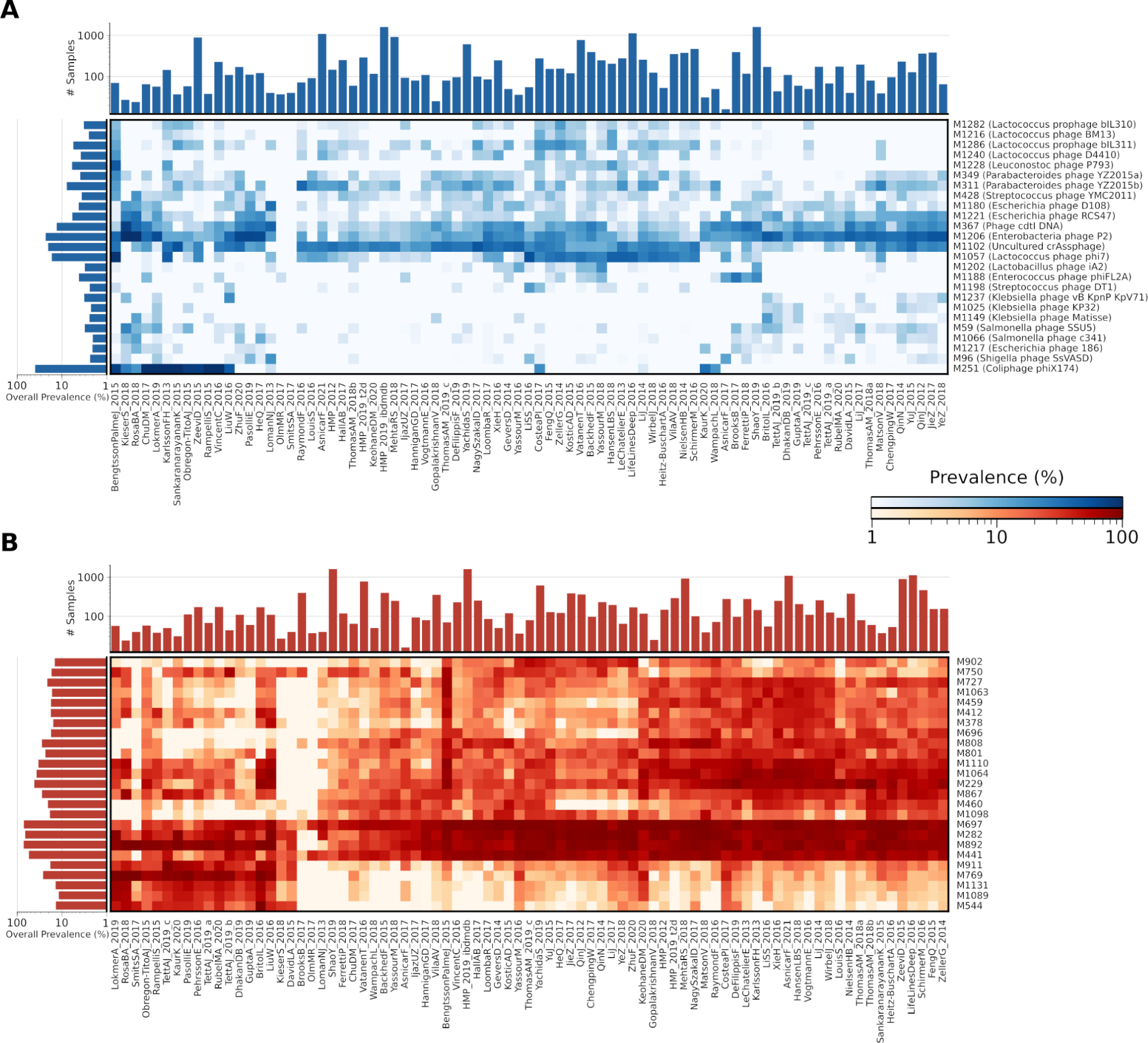
Prevalence of the 1,345 VSGs in 81 datasets and 18,756 human gut metagenomes. Per-dataset prevalence of the top 100 known (**A**) and unknown (**B**) Viral Sequence Groups (VSGs, rows) in 18,756 samples of 81 datasets (columns). The barplot on top of the heatmap reports the number of samples in each dataset. The barplot on the left indicates the average prevalence of each VSG across all datasets (when the VSG is detected at least once). Values are reported in **Supplementary Table 7**.

Bacteriophages can alter the microbial communities and change the relative abundance of microbes susceptible to infection. Among the 1,345 VSGs included in MetaPhlAn 4.1, 90.6% had at least one hit with a microbial species (**Fig. 3C**), while as many as 12% groups were associated with more than 30 microbial species. This remained true even considering individual viral sequences instead of groups, with 78.4% of the 45,872 viral sequences with at least one species (**Supplementary Fig. 4A**). This finding confirms that bacteriophages may have a broad species tropism, as suggested in other studies ^64^. We then used these associations to compare the prevalence of microbes in 18,714 gut metagenomes with and without their associated viral groups. We found higher microbial prevalence when the viral groups were co-present with their target species, both with known and unknown VSGs (**Fig. 6**). Overall, these results highlight how novel viral genome clusters incorporated into MetaPhlAn 4 can be efficiently detected from metagenomes, yielding a combined prokaryotic and viral composition to better characterize the diversity, ecology and host-condition association of microbiomes.

**Figure 6.**
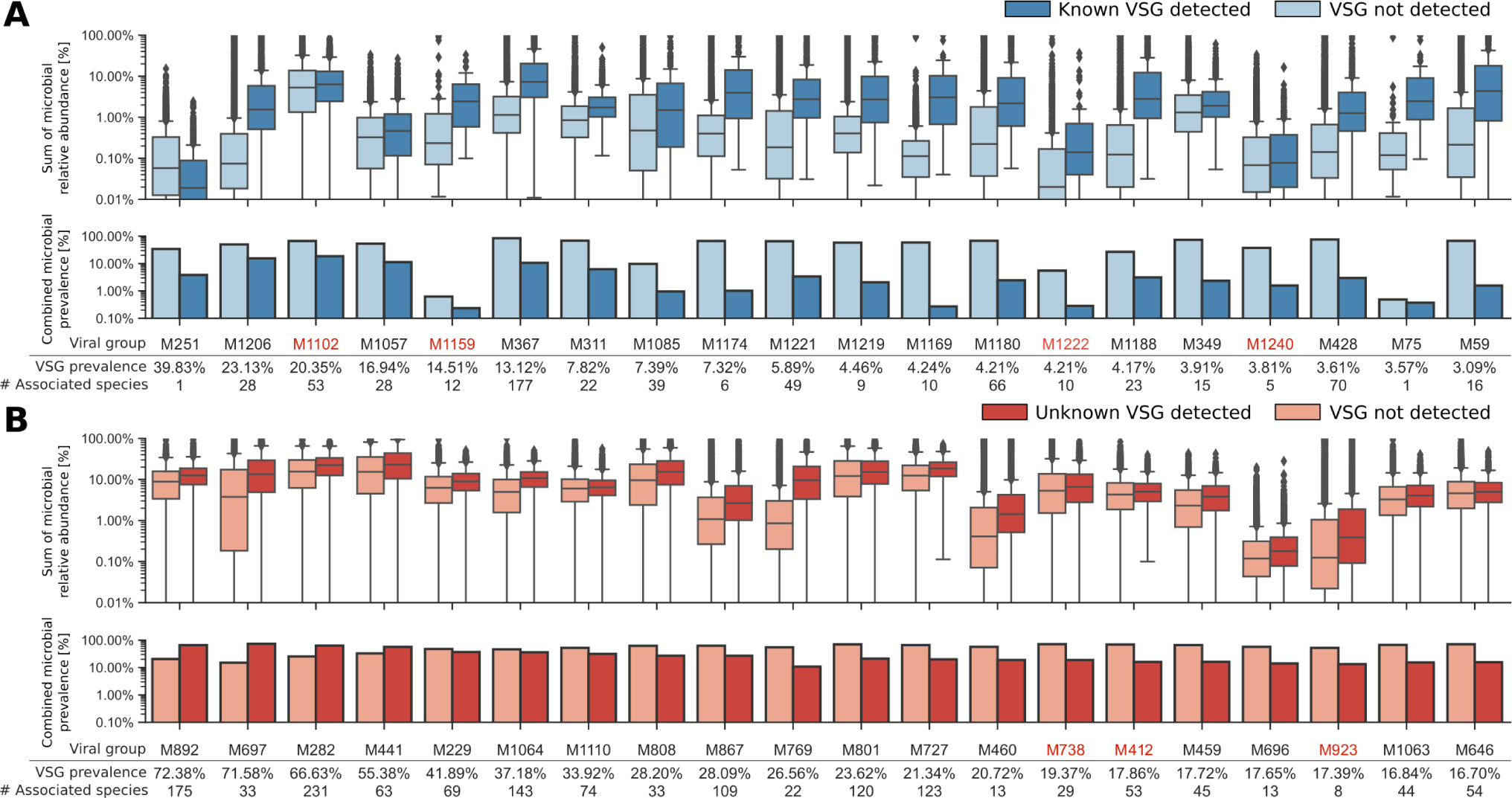
Abundance of microbial species associated with bacteriophages, with and without co-infection. **(A-B)** CRISPR spacers were used to predict the associated microbial species for the most prevalent known (**A**) and unknown (**B**) Viral Sequence Groups (VSGs). The relative abundances of microbial species associated with each VSG were calculated using a custom version of the MetaPhlAn taxonomy classifier that also profiles Species-level Genomes Bins (SGBs). The boxplots show the relative abundances of SGBs associated with the Viral Groups in 18,714 samples, with (dark) or without (light) the bacteriophage. If the Viral Group is associated with more than one species, the sum of the relative abundances for each associated species in each sample is shown. The barplot under the boxes reflects the prevalence of all the SGBs associated with a Viral Group, while the number of SGBs associated with each group and the group average prevalence in metagenomes are reported below the bar. Boxes encompass distribution quartiles, and whiskers extend to 1.5 IQR. Groups labeled in red are those where no statistical significance was found when comparing the microbial abundances with or without viral presence (Mann-Whitney U test, p >0.05). Viral Groups undetected in at least two datasets were excluded. Host-phage associations are reported in **Supplementary Table 8.**

## Discussion

We presented here a novel strategy that exploits Viral Like Particle (VLP) enriched viromes as the first crucial step in a multiple-step pipeline for the identification and discovery of potentially new viral sequences. Compared to current methods to label as viral contigs of metagenomic origin ^19,24,74^, our approach maximizes the chances of identifying phages with no common characteristics in terms of sequence, structure, or derived features with known phages. This is derived from our specific focus on the metagenomic assemblies from carefully evaluated highly enriched viromes, under the assumption that they would yield more sequences of sure viral origin. We accordingly found that almost 20% of the 5,651 HEVCs were never identified as viral by any of the four viral-prediction tools we tested, and 39.1% were labeled as viral by one tool only (**Fig. 2**). Fully *de novo* identification of phages via selection of maximally enriched virome is thus an effective approach that can be further exploited to continue unraveling genomic viral dark matter.

Expansion of the novel unraveled viral diversity using massive sets of metagenomes highlighted several aspects of gut phage communities. First, we were able to reconstruct phylogenies of some of the most prevalent viral clusters, including *Lactococcus* and *Klebsiella* phages, as well as the recently characterized crAssphage ^61^. Second, several prevalent unknown viral clusters were extremely conserved, with pairwise sequence identities as high as 99%, and such high-identity within-cluster sequences were also shared across different studies and individuals. Third, for several known phages we expanded the phylogenetic structure with relevant clades: for example, when the original 13 crAssphage HEVC sequences were used to find homologous contigs in viromes and metagenomes, the 277 new sequences formed a separate phylogenetic group that hence expanded the *crAssphages* diversity. Continued analysis of phage databases can substantially deepen these aspects and move the analysis of phages closer in resolution to that available for bacterial members of microbiomes.

We comprehensively analyzed our novel resource to link phages with their bacterial hosts. Matching phages to their host(s) via CRISPR spacers is not only highly effective in its primary goal (76.21% of the 1,345 viral groups matched to a species at least three spacers and only 9.37% had no associations), but also provides further validation of the phagic nature of the identified sequence (78.42% sequences and 88.24% of the viral clusters matched with at least one CRISPR spacer) and enables ecological modeling of phage-host interactions. On the latter, we found that phages tend to co-occur with their hosts in microbiomes (**Figure 6**) pointing at more complex and larger types of phage-bacteria interactions in which symbioses may play a more important role than previously appreciated.

Several aspects need further investigations and follow-up studies. Some sequences within clusters matched partially against known plasmids. While a viral origin of such elements cannot be excluded, the fact that plasmid sequences may still be present in the highly-enriched viral contigs poses the need for extra caution in the interpretation of the results when such extremely abundant sequences are found. To overcome this limitation, further research is needed to characterize the nature of the sequences contained in our collection. Moreover, some of the sequences we retrieved might align to prophagic regions in microbial genomes; while CRISPR spacers have been extensively used to pinpoint viral-host interactions, and it is unlikely that sequences highly recognized by CRISPR-Cas9 systems can frequently be prophages, complementary analyses may be needed to carefully disambiguate sequences coming from integrated or non-integrated phages. Finally, our work largely focused on human gut microbiomes, but similar studies should be performed on several other host-associated or environmental microbiome types.

Importantly, the large-scale *de novo* analysis of phage diversity we performed here cannot be readily applied to novel viromes and metagenomes without a high-performance computing system and advanced computational expertise. To this end, in addition to the curated catalog of viral sequences we provide publicly (see **Data Availability**), we also incorporated the database within MetaPhlAn 4.1, allowing it to detect these new viral taxa in metagenomes. Currently, detection of viruses is carried out by read-to-genome matching filtered by breadth of coverage, leaving opportunities for the development of new methods for more specific profiling (e.g. based on marker sequences or k-mer matching). For further expanding on the purposes of profiling, the database can also be more extensively integrated with others (including viruses from outside the human gut), and additional approaches can be explored to capture the extreme variability of viral taxonomy and phylogenetic divergence. However, any such methods require the extensive catalogs of new viral diversity as developed here, which can then be applied for efficient metagenomic profiling in the human microbiome and beyond.

## Materials and Methods

Our pipeline for viral discovery exploits highly enriched viromes that undergo metagenomic assembly to produce potentially viral contigs. These contigs are then screened to remove unwanted residual contaminants (i.e., microbial non-viral sequences). Finally, contigs are clustered by similarity, and clusters are then extended by retrieving thousands of sequences from assembled metagenomes and lowly enriched viromes. In the following, we will detail the methodological aspects of our pipeline, and its application to 3,044 viromes and 9,721 metagenomes to retrieve the 3,944 viral sequence clusters (VSCs). The pipeline is depicted in **Fig. 7** and **Supplementary Fig. 7**.

**Figure 7.**
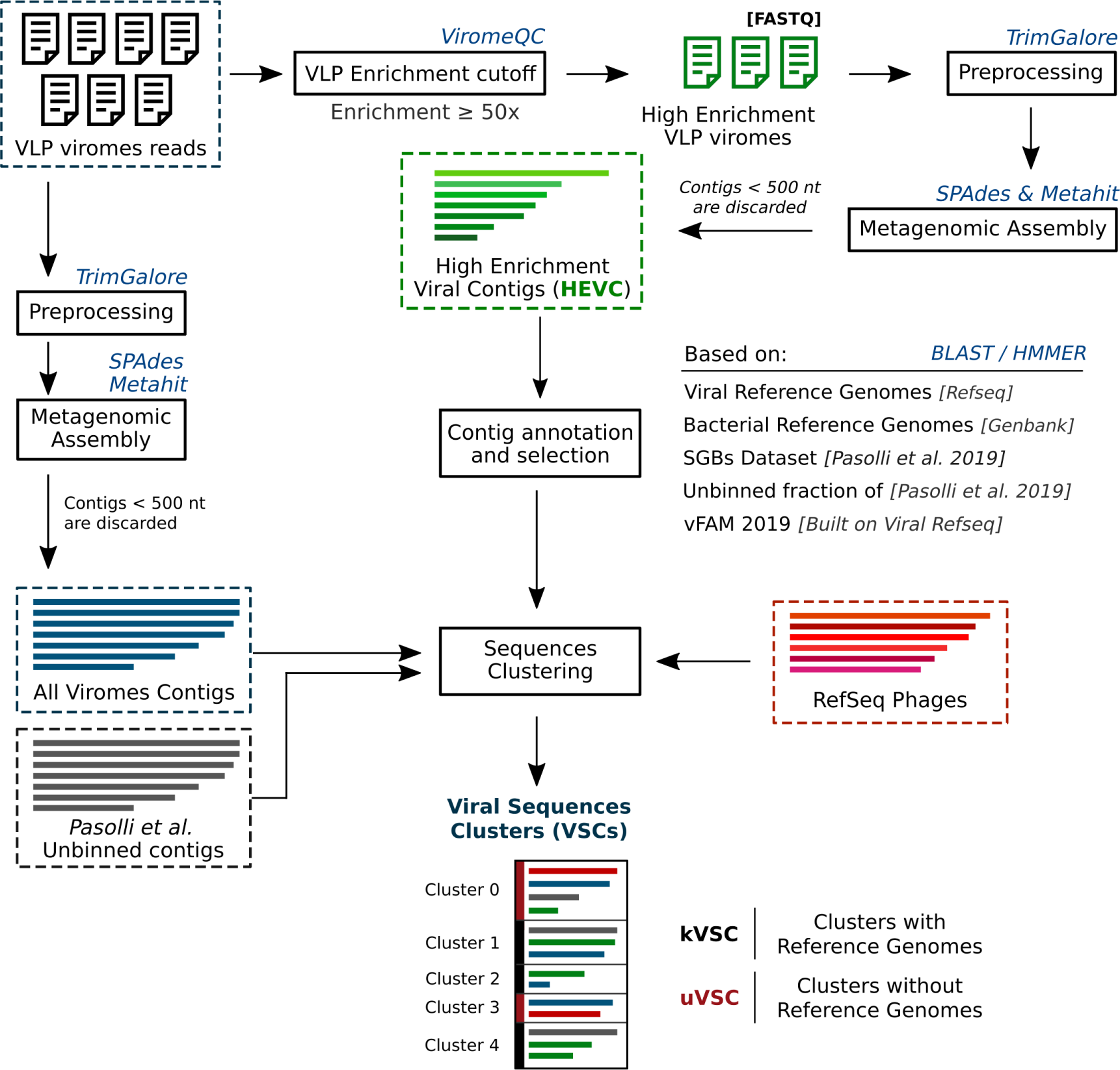
Overview of the pipeline for the assembly and identification of viral contigs. The flowchart represents the steps followed to select highly enriched viral contigs from viromes, and to produce clusters of viral sequences integrating virome and metagenomic contigs. Viromes were preprocessed, assembled, and screened for enrichment with ViromeQC. Contigs from samples with an enrichment score higher than 50x were labeled as Highly Enriched Viral Contigs (HEVCs). HEVCs were screened to reduce further sources of non-viral contamination by mapping against reference bacterial genomes and Metagenomic Assembly Genomes (MAGs). The remaining HEVCs were clustered with a) reference viral genomes and b) sequences similar to HEVCs retrieved from unenriched metagenomes and viromes. This clustering step produced the set of Viral Sequence Clusters (VSCs), which were split into known (kVSCs, if they contained at least one reference viral genome) and unknown (uVSCs, if they did not contain any reference viral genome). A detailed representation of the Sequence Clustering procedure is shown in **Supplementary Fig. 7**.

### Retrieval of metagenomes and viromes used in the analysis

A total of 3,044 viromes were retrieved from NCBI-SRA with SRA-toolkit. This includes the viromes analyzed for viral-enrichment in the scope of a previous work ^39^ and viromes from 14 additional datasets. All accession numbers are reported in **Supplementary Table 1**. Samples that were replicated by more than one NCBI-SRA run were downloaded and processed independently. Prior to metagenomic assembly, viromes were preprocessed with Trim Galore version 0.4.4 ^75^ to remove low-quality (i.e., Phred quality < 20) and short (i.e., read length < 75) reads (parameters: --stringency 5 --length 75 --quality 20 --max_n 2 --trim-n). Reads aligning to the human genome *hg19* were removed by mapping with Bowtie2 version 2.4.1 ^76^ in “end-to-end” global mode. The preprocessed reads were kept ordered and split in forward, reverse, and singletons FASTQ files, when possible (i.e., for paired-end sequencing libraries). In total, 2.76×10^10^ high-quality reads were retrieved.

Publicly available datasets were used to annotate the virome-contigs (see below, contig annotation). In particular, sequences of the metagenome-assembled genomes (MAGs) from Pasolli *et al.* ^32^ were retrieved from http://segatalab.cibio.unitn.it/data/Pasolli_et_al.html, together with the assembled contigs that could not be binned by MetaBat2 (i.e., the unbinned fraction). Five additional metagenomic datasets (*CM_ethiopia*, *Heitz-BuschartA_2016*, *KieserS_2018*, *RosaBA_2018*, and *ShiB_2015*, see **Supplementary Table 1** of Pasolli *et al.*) were binned as described in the original study but were not grouped into Species-level Genome Bins (SGBs). MAGs and unbinned contigs from three additional metagenomic datasets from Tanzania, Ghana, and Madagascar were also included ^77^. For cases in which the SGB classification lacked, we interpreted each bin as an individual SGB for annotation purposes. In this paper, contigs that are neither assigned nor binned into any SGB are referred to as “unbinned contigs” or “unbinned fraction”.

### Virome Enrichment Estimation

The viral enrichment of the VLP-viromes was assessed with ViromeQC v. 1.0 ^39^, a computational tool that can be applied to raw metagenomic reads to estimate the efficacy of VLP enrichment. ViromeQC calculates the percentage of reads aligning against two sets of markers of sources of microbial contamination: a) the small and large subunits of the 16S/18S and 23S/28S rRNA genes, and b) a set of single-copy bacterial markers. It then outputs an enrichment score calculated on the abundances of the microbial markers. A score lower than 1x indicates negative enrichment (i.e., the sample is less enriched than a metagenome). Samples with a ViromeQC score of 50x or higher were considered to be highly enriched (**Table 1**). RNA-viromes and samples where RNA and DNA were co-extracted were not considered for the viral enrichment estimation.

### Metagenomic Assembly

After preprocessing, all the 3,044 viromes underwent metagenomic assembly. Paired-end samples were processed with custom Python scripts (see **Code Availability**) to preserve the “read pairing” information in the FASTQ format and were then assembled with metaSPAdes version 3.10.1 ^78^ (k-mer sizes: -k 21,33,55,77,99,127). Unpaired samples were assembled with Megahit version 1.1.1. (default parameters). Contigs shorter than 500 nucleotides were discarded, and 4.11 x 10^7^ contigs were retained.

### Contigs Annotation

To focus on human-gut-associated viruses, only contigs originating from a gut virome (120,041 contigs, see **Supplementary Table 2**) were further analyzed. Open Reading Frames (ORFs) were predicted using Prokka version 1.12 ^79^, with --kingdom Viruses. To determine the similarity of the contigs to bacterial and viral reference genomes, contigs were mapped against a) all the reference viral genomes in RefSeq ^41^, release 91; b) a set of 80,853 complete and draft bacterial genomes from NCBI GenBank ^32,80^; c) the MAGs and reference genomes from Pasolli *et al.* All mappings were performed with BLAST, version 2.6.0 ^81^ in nucleotides space and with the default parameters except for -max_target_seqs 1000. BLAST databases were built with custom scripts to allow for better hits-tracking in the downstream analysis. Only hits with an alignment length >=1000 nucleotides and a percentage of identity >=80% were considered. We also computed the total breadth-of-coverage of each contig against each database (i.e., the bacterial genomes, viral genomes, and SGBs), by calculating how many nucleotides of the contig were covered by at least one blast hit in the database. Values for each contig are reported in **Supplementary Table 2**.

Contigs were also mapped against the unbinned fraction of Pasolli *et al.* to identify potentially viral sequences in the residual unbinned fraction of binned MAGs. In this case, we set -max_target_seqs 500000 to maximize the number of hits to unbinned contigs. Only hits with length >=1000 nucleotides and identity >=80% were considered. We calculated the proportion of ORFs that matched viral proteins in each contig, ORFs were mapped with hmmscan version 3.1b2 ^82^ against a database built with the V-Fam code provided by Skewes-Cox *et al.* ^68^. We used all the proteins from Viral Refseq (release 91). An ORF was considered to match a viral protein domain if it had at least one full-domain hit with an e-value lower than 10^-5^.

To determine whether a contig was present in more than one sample or dataset, we ran an all-vs-all BLAST search on the 120,041 sequences. To be detected in another sample, one contig needed to have an overlap of at least 1000 nucleotides with at least 80% identity. The number of hits to other viromes is reported in **Supplementary Table 2** and is referred only to the 255 highly enriched viromes shown in **Table 1**.

### Retrieval of 2,619 bacteriophages from RefSeq

We downloaded all the viral genomes from RefSeq ^41^ (Release 99; n=12,182) and selected the phages in two steps. First, we used the information present in the sequence ID to retrieve all the sequences containing the keyword “phage” (ignoring case distinctions) in the header (n=2,543); then, we relied on the NCBI taxonomy (taxid = 10239) to refine our selection, adding 76 more sequences to our set.

### Selection of the 5,651 High-Enrichment Viral Contigs

To further remove contigs likely part of bacterial or archaeal genomes, we used the Pasolli *et al.* MAGs and metagenomes dataset to discard sequences that a) were found binned in the same Species-level Genome Bin (SGB) in more than 30 metagenomes; or b) were found binned in any SGB in more than 50 metagenomes. Finally, only contigs longer than 1,500 nucleotides and that were found in the unbinned fraction in more than 20 metagenomes were kept. Of the initial 120,041 contigs, 4,952 met the selection criteria. These contigs were added to the viral reference genomes of bacteriophages that could be found in at least 20 metagenomes (n=699). In total, 5,651 sequences were then considered as “High-Enrichment Viral Contigs” (HEVC).

### Analysis of the reconstructed contigs with viral-prediction software

Viral prediction tools were used to classify the retrieved contigs as viral or non-viral. Specifically, VirSorter version 1.0.5 ^20^ was run using diamond ^83^ (--diamond) with the RefSeq database (--db 1), both in standard and decontamination mode (--virome parameter). Predictions falling in VirSorter classes 1,2,4 and 5 were considered as viral. VirFinder version 1.1 ^19^ was run in R 3.6.3 using the standard prediction model, a threshold score >=0.8 and a p-value lower than 0.05. ViralVerify version 1.0 was run with the provided database of virus-chromosome-specific HMMs ^21^. Seeker version 1.0.1 ^23^ was run with the default parameters and contigs with a score >=0.8 were considered predicted as viral.

### Clustering of the 5,651 High-Enrichment Viral Contigs and clusters extension

The 5,651 sequences from highly enriched viral contigs and reference genomes were clustered into 3,944 clusters at 90% identity with VSEARCH version 2.14.2 ^59^, (parameters --cluster_fast --id 0.9 --strand both --maxseqlength 200000). Clusters that contained at least one RefSeq viral genome were labeled as known viral sequence clusters (kVSC); clusters that only contained novel sequences were labeled as unknown (uVSC).

Clusters were extended with contigs from viromes and metagenomes. We took contigs from the full dataset of 3,044 assembled viromes (i.e., highly enriched and non-highly enriched viromes, **Supplementary Table 1**) and from the unbinned fraction of metagenomes analyzed in Pasolli *et al.* (see “Selection of the 5,651 High-Enrichment Viral Contigs”). First, these contigs were mapped against the 5,651 HEVCs with BLAST ^81^. Then, sequences with a percentage of identity of at least 80% over at least 1000 nucleotides to an HEVC were kept (69,484 contigs similar to an HEVC from viromes, 355,825 from unbinned metagenomes). The 425,309 contigs were used to build a sketch database with Mash version 2.0 ^62^ (command: mash sketch -i -s 10000). Next, each VSC centroid (i.e., the longest sequence within each of the 3,944 initial clusters) was mapped against the initial set of contigs using Mash (command: mash dist -d 0.1 -v 0.05), and contigs with a distance lower than 10% (p-value <=0.05) were assigned to the closest VSC cluster (i.e., the VSC with the minimum mash distance), hence extending clusters with new sequences and producing extended-HEVCs.

Extended-HEVCs were then further re-clustered at 70% identity with VSEARCH, and only clusters that contained at least one of the original 5,651 sequences were kept. Clusters that had more than one valid sub-cluster were kept separated. For example, cluster *vsearch_c1003* contained five sub-clusters, two of which contained sequences from the starting 5,651 HEVCs. Hence, two sub-clusters of *vsearch_c1003* (c1003_c0 and c1003_c3, see **Supplementary Table 3**) were kept. This step produced the final 4,077 extended clusters.

### Cluster representatives selection and prevalence analysis

We built a collection of representative sequences for each of our 3,944 VSCs to select sequences that maximized the diversity within each cluster. Hence, we re-clustered all the sequences within each VSC at 95% identity with VSEARCH (parameters: --cluster_fast --id 0.95 --strand both). We then selected as final representatives: a) the centroids of the 95% clustering; b) all the original 5,651 contigs from highly-enriched samples and reference genomes. In total, 47,820 clusters-representative sequences were selected (median length: 6,355 bp; maximum length: 240 kbp). Sequences were de-replicated at 99% identity and 90% overlap with CD-HIT version 4.6.8 ^84^ (parameters -n 10 -d 0 -c 0.99 -aL 0.9). The final collection of 45,872 sequences is publicly available together with cluster metadata (see **Data Availability**).

The 45,872 sequences were further grouped into 1,345 Viral Sequence Groups (VSGs), by taking the connected components of a graph built as follows. A pair of vertices (sequences of different VSC clusters) were connected if the longest sequence was covered for at least 33% of its length (2000nt minimum) by alignments of the shortest sequence (avg. percentage of identity >=90%). Only alignments longer than 500nt were considered to calculate the combined pairwise coverage.

### VSCs prevalence in metagenomes

To calculate the prevalence of each VSG in metagenomes, we mapped the raw reads of 18,756 publicly available metagenomes against the representative sequences of each VSC. Mapping was performed with Bowtie2 version 2.4.1 ^76^, in end-to-end global mode. The aligning reads were then re-mapped agains the best target within each VSG. The best target was defined as the sequence with the highest number of alignments, normalized by the length of each sequence. Alignments were converted into BAM files with Samtools version 1.3.1 ^85^. Breadth and depth of coverage for each cluster were calculated with the PySam package. The minimum breadth of coverage to consider a cluster as detected was set to 75%.

### Phylogenetic analysis, data analysis, and visualization

To build phylogenetic trees, we selected the sequences with a length within 25% from the median length of all the sequences in each cluster. Sequences belonging to the original 5,651 HEVCs were kept regardless of their length, unless differently specified. Multiple-sequence-alignments for elements of each Viral Sequences Cluster were performed with MAFFT version 7.453 ^86^, with automatic parameters selection (--auto). Alignments were trimmed with Trimal version 1.2rev59 ^87^, with the -gappyout option. Phylogenetic trees were computed using RAxML version 8.1.15 ^88^ with the GTRGAMMA model and bootstrap analysis (parameters: -p 48315 -x 48315 -# 100 -m GTRGAMMA -f a). We performed a manual curation and topology-checking of the resulting phylogenetic trees, branch lengths, and bootstrap values. In some cases, we also took into consideration the corresponding alignment to identify possible sources of non-phylogenetic signal. Trees with very low bootstrap values (below 40 for all nodes), excessively long and isolated branches (i.e., showing a number of substitutions per site more than 10 times the median branch length for that specific tree and containing clades with fewer than 4 sequences), and/or fragmented alignments (cases in which half or more of the alignment consisted of short regions of less than 20 bp with similarity below 30%) were discarded.

Pairwise sequence identities of each viral sequence cluster were calculated on the multiple-sequence alignments. To ensure that only sequences of similar lengths were aligned, we further filtered the alignments for each cluster by retaining only sequences within 25% of the median sequence length of the HEVCs in the cluster. The sequence identity of two sequences was defined as 1-Hamming_Distance (seq1,seq2), without considering gaps.

Hierarchical clustered heatmaps were generated with hclust2 ^89^ using Bray-Curtis distance to cluster features (VSCs) and correlation distance to cluster samples. Average linkage was used in the hierarchical clustering. Plots and figures were drawn with matplotlib ^90^ and Seaborn ^91^. Upset plots were drawn in Python with the upsetplot package, version 0.4 ^92^. Bioinformatic analysis on sequences was performed in Python3 with packages BioPython ^93^, Pandas version 1.0.1 ^94^, and NumPy version 1.18.1 ^95^. Computation was performed on the High-Performance Computing infrastructure of the University of Trento, Italy.

### CRISPR Extraction and Phage-Host associations

CRISPR spacers were predicted from 377,346 metagenomic assembled genomes (MAGs) and 169,111 reference genomes from the Jan21 release of the ChohoPhlAn/MetaRef database ^40^ using MinCED with default settings, version 0.4.2, derived from CRT ^96^. Spacers with more than 33% or 25 undetermined nucleotides (Ns), were excluded. Spacers were de-duplicated with CD-HIT version 4.6.8 ^84^, and mapped with BLAST version 2.6.0 ^81^ (parameters: -word_size 7 -max_target_seqs 500000 -evalue 0.01) against the 45,872 viral sequences presented in this work, and the 142,809, 33,242, and 2,377,994 sequences in GPD ^64^, GVD ^97^, and IMG/VR 2020 ^65^, respectively. To associate viruses and hosts, at least one CRISPR spacer (query) had to map at <3 SNPs of distance to the target (viral sequence), with a query coverage >90% and a percentage of identity >90%.

### Code and Data Availability

The code used to perform this analysis is available on GitHub at https://github.com/SegataLab/viromedb. The viromes used in this study are described in the ViromeQC publication ^39^. Additionally, viromes from 14 other studies were used (**Supplementary Table 1**). The MAGs used to annotate viral contigs are described in Pasolli *et al.* and at http://segatalab.cibio.unitn.it/data/Pasolli_et_al.html. The metagenomes used to extend VSCs are referred to in the same publication. The representative sequences of all the VSCs are available on Zenodo (DOI: http://doi.org/10.5281/zenodo.10512460), and at http://segatalab.cibio.unitn.it/data/VDB_Zolfo_et_al.html. The module to profile viral sequence groups in raw metagenomes is available in MetaPhlAn 4.1 ^40^ using the --profile_vscs option.

## Acknowledgments

We thank all the members of the Segata and Huttenhower lab for fruitful discussions on this work. This work was supported by the European Research Council (ERC-STG project MetaPG-716575 and ERC-CoG microTOUCH-101045015) to N.S., by European Union’s Horizon 2020 research and innovation programme (ONCOBIOME-825410 project, MASTER-818368 project, and IHMCSA-964590) to N.S., by the MUR PNRR project INEST-Interconnected Nord-Est Innovation Ecosystem (ECS00000043) funded by the NextGenerationEU to N.S., by the National Cancer Institute of the National Institutes of Health (1U01CA230551) to N.S., by the Premio Internazionale Lombardia e Ricerca 2019 to N.S., by the JST AIP Acceleration Research (JPMJCR19U3) to T.Y, and by the Japan Society for the Promotion of Science (KAKENHI JP16H06279 (PAGS)) to T.Y. C.M. is funded by the Chronic Disease Research Foundation. Support for this work was also provided by UKRI/MRC grants MR/W026813/1 and MR/Y010175/1 to C.M. A part of this research is supported by the COI-Next Program of JST/MEXT to OIST.

