## Supplementary Figures and Tables for "Discovering and exploring the hidden diversity of human gut viruses using highly enriched virome samples"

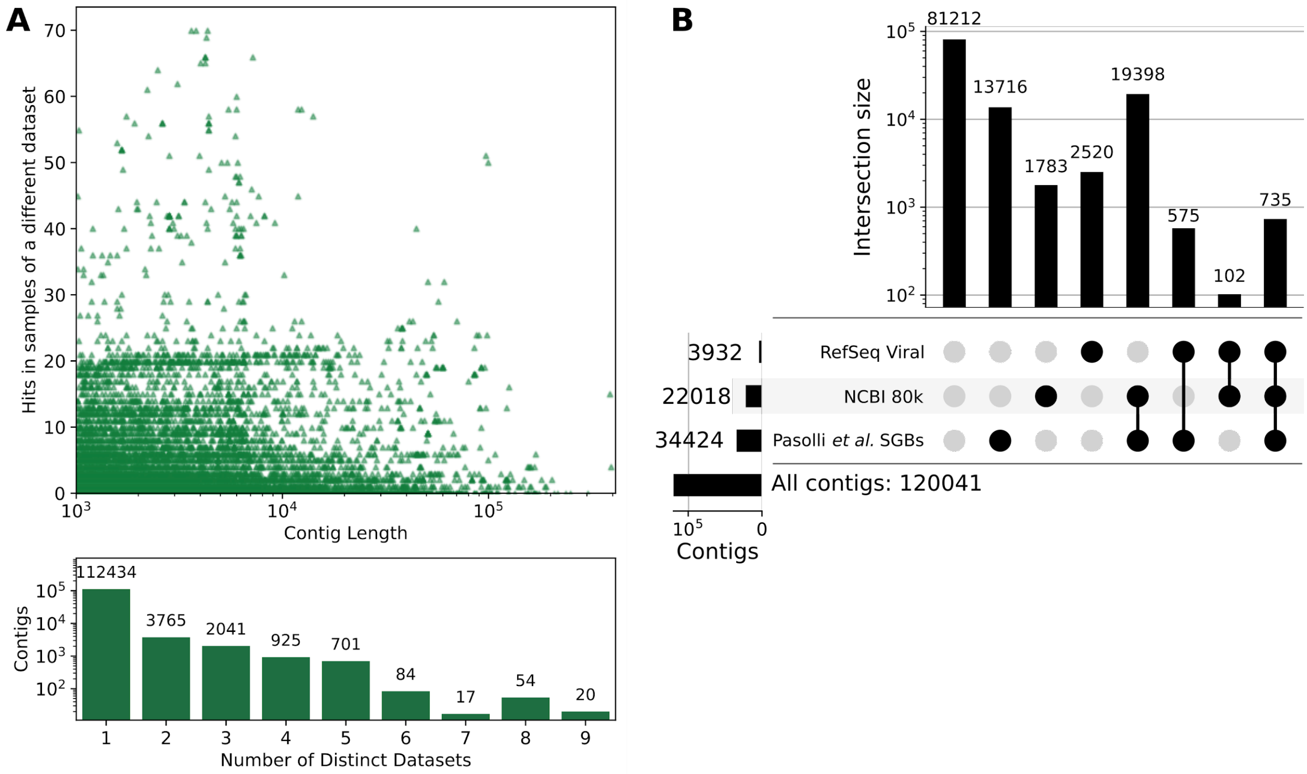


**Supplementary Figure 1. Origin and prevalence of the 120,041 contigs assembled from highly-enriched viromes. (A)** Prevalence of contigs across 11 virome studies**.** The scatterplot shows each of the 120,041 contigs assembled from the 255 highly enriched gut viromes. X-axis represents contig length, while the y-axis shows the number of hits in samples of a different dataset. The bottom bar plot indicates the number of contigs found in a given number of distinct datasets. Detection of a contig in multiple samples was based on BLAST hits longer than 1000 nucleotides and with a percentage of identity of 80% or higher (see **Methods**). **(B) Detection of the 120,041 contigs assembled from highly enriched gut viromes in other sequence resources.** The upset plot shows the intersection size of contigs covered by hits against the RefSeq viral references, a set of bacterial reference genomes (NCBI 80k), and the MAGs from Pasolli et al., 2019. Contigs are annotated in each set if they had a cumulative breadth of coverage >50% with any reference sequence in the set. Intersection sets are highlighted by the continuous vertical lines in the bottom part of the plot.

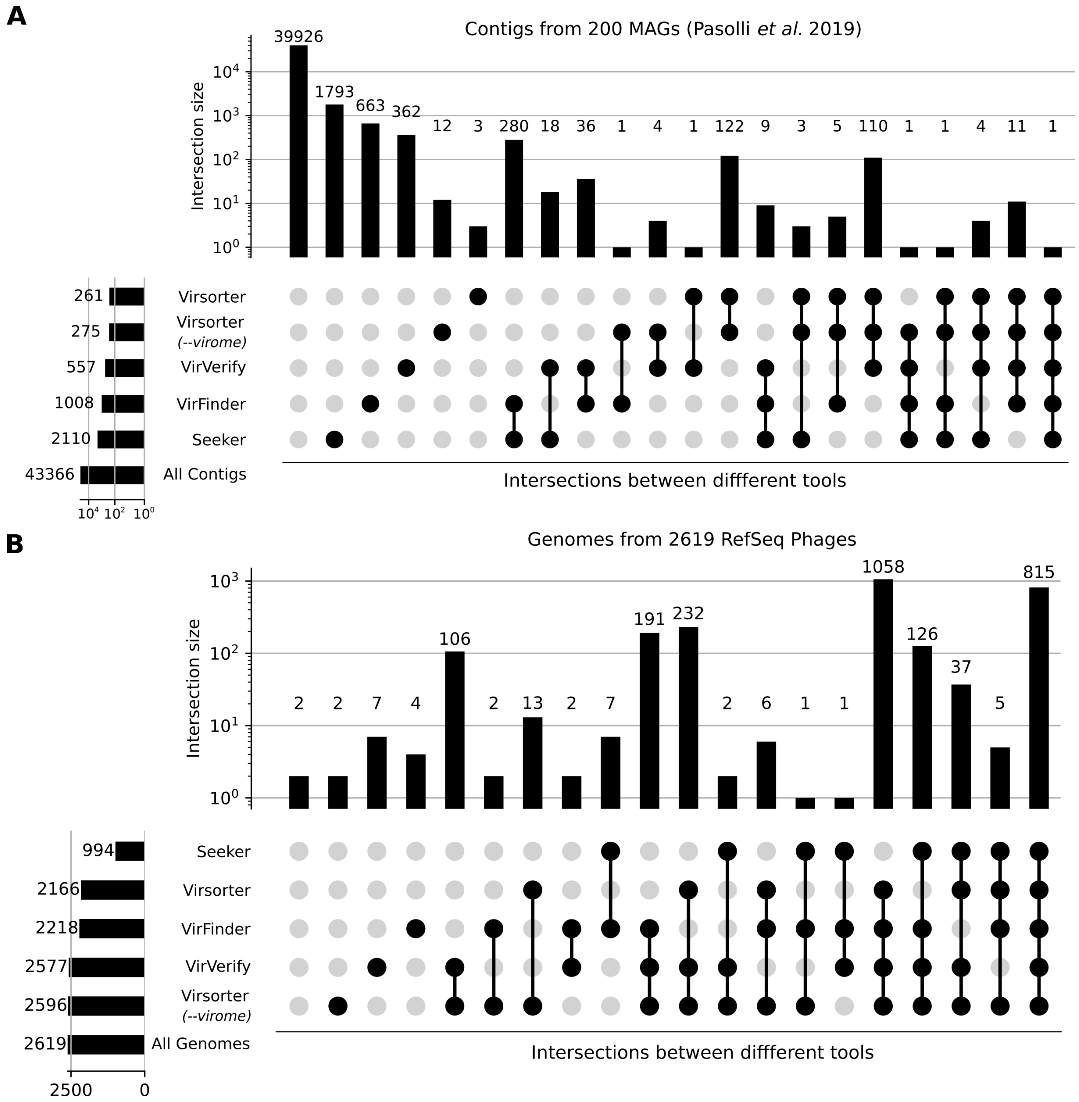


**Supplementary Figure 2. Comparison of different Viral Detection tools on Viral and Bacterial Genomes.** Four viral-detection tools were applied to a set of contigs from medium and high-quality MAGs (**A**) and to 2,619 reference genomes of bacteriophages (**B**). The plot represents the number of contigs classified in each intersection set. 'All genomes' and 'All contigs' refer to the full set of analyzed contigs and genomes.

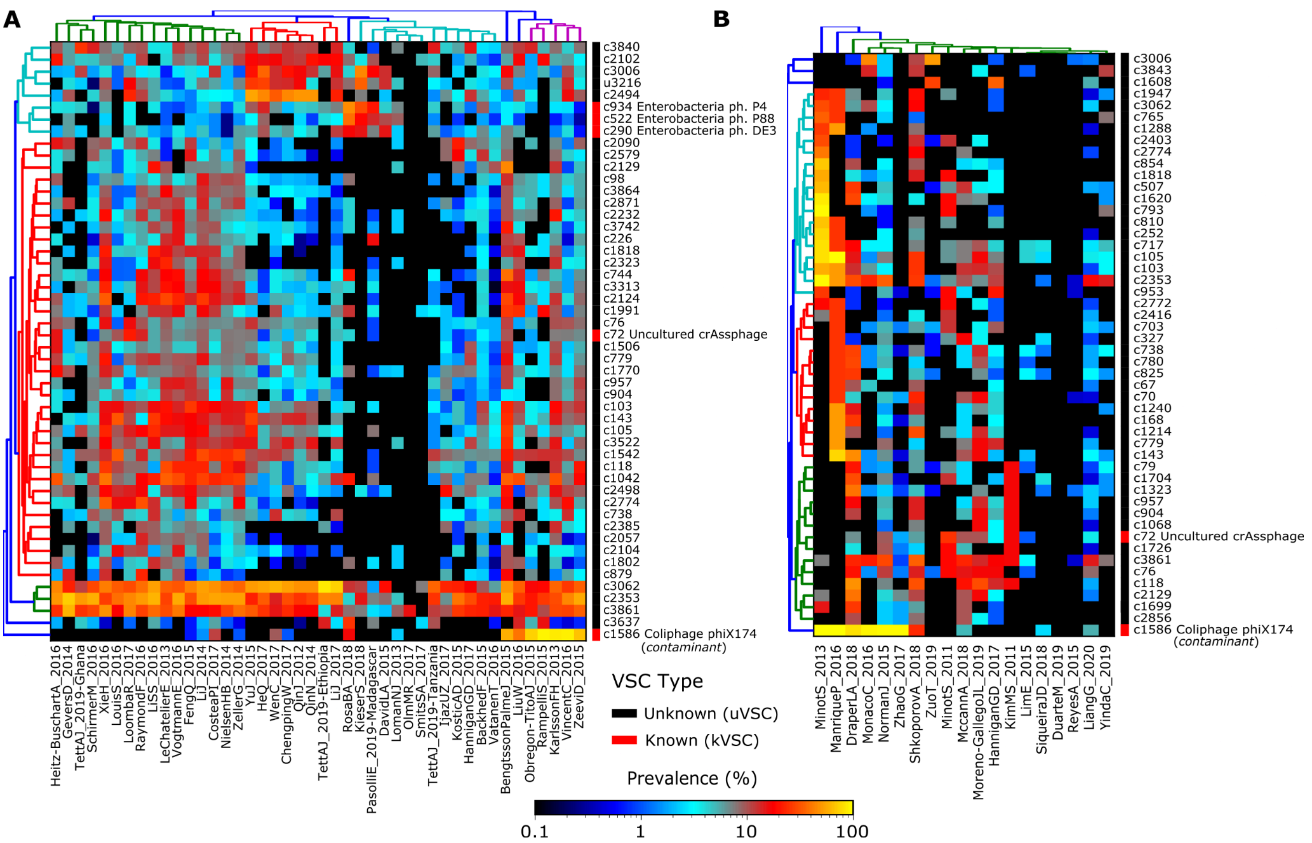


**Supplementary Figure 3. Prevalence of the Viral Sequence Clusters (VSCs) in Metagenomes and Viromes.** The clustered heat maps show the number of detected VSC across gut metagenomes **(A)** and viromes **(C)** normalized by the number of samples in each dataset. Right-side boxes indicate known (kVSC, red) and unknown (uVSCs, black) clusters. Known clusters are labeled with the name of the closest reference. Detection of VSCs is based on BLAST alignments against assembled contigs from metagenomes and viromes.

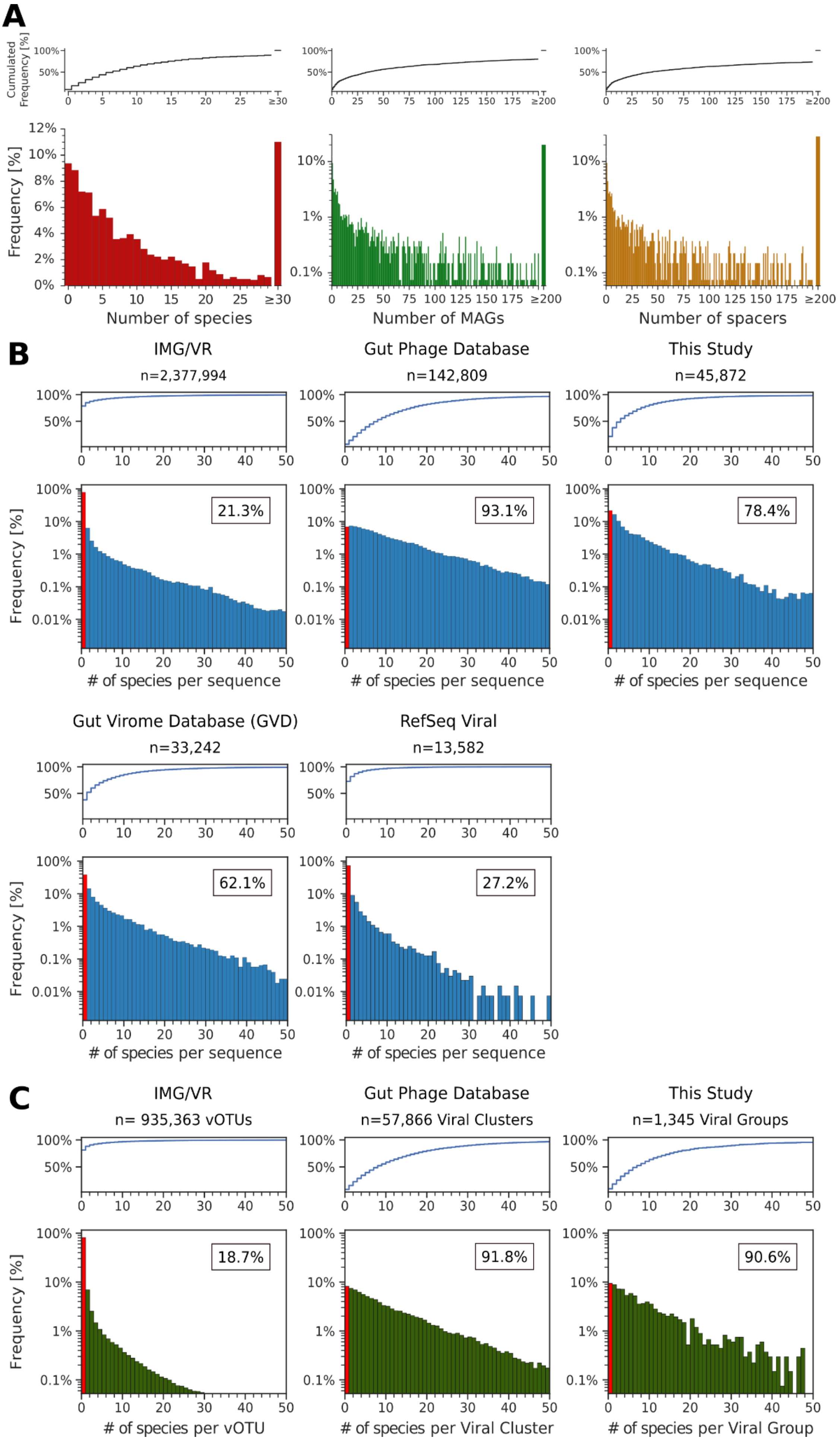
A
Cumulated Frequency [%]
Frequency [%]
Number of species
Number of MAGs
Number of spacers
B
IMG/VR
n=2,377,994
21.3%
Gut Phage Database
n=142,809
93.1%
This Study
n=45,872
78.4%
Gut Virome Database (GVD)
n=33,242
62.1%
RefSeq Viral
n=13,582
27.2%
### of species per sequence
C
IMG/VR
n= 935,363 vOTUs
18.7%
Gut Phage Database
n=57,866 Viral Clusters
91.8%
This Study
n=1,345 Viral Groups
90.6%
### of species per vOTU
### of species per Viral Cluster
### of species per Viral Group

**Supplementary Figure 4. CRISPR Spacers distribution in multiple viral catalogues. (A)** Percentage of the 1,345 viral groups (y-axis) that are associated with a given number (x-axis) of distinct microbial species (red left histogram), MAGs (green middle histogram) or CRISPR-spacers (orange right histogram). The cumulative distribution is reported above each histogram. CRISPR spacers were predicted from 377,346 Metagenomic Assembled Contigs and reference genomes (see Methods). Viral groups and microbial hosts were considered associated if the predicted microbial CRISPR-spacer could be detected in a sequence within the viral group. **(B-C)** Percentage of the viral groups and sequences in different viral catalogues that are associated with a given number of distinct microbial species. Red bars indicate the percentage of sequences that were not hit by any CRISPR spacer. The inset number is the percentage of sequences assignable to a host by CRISPR spacers. Frequencies are normalized by the number of sequences **(B)** or clusters **(C)** in each study.

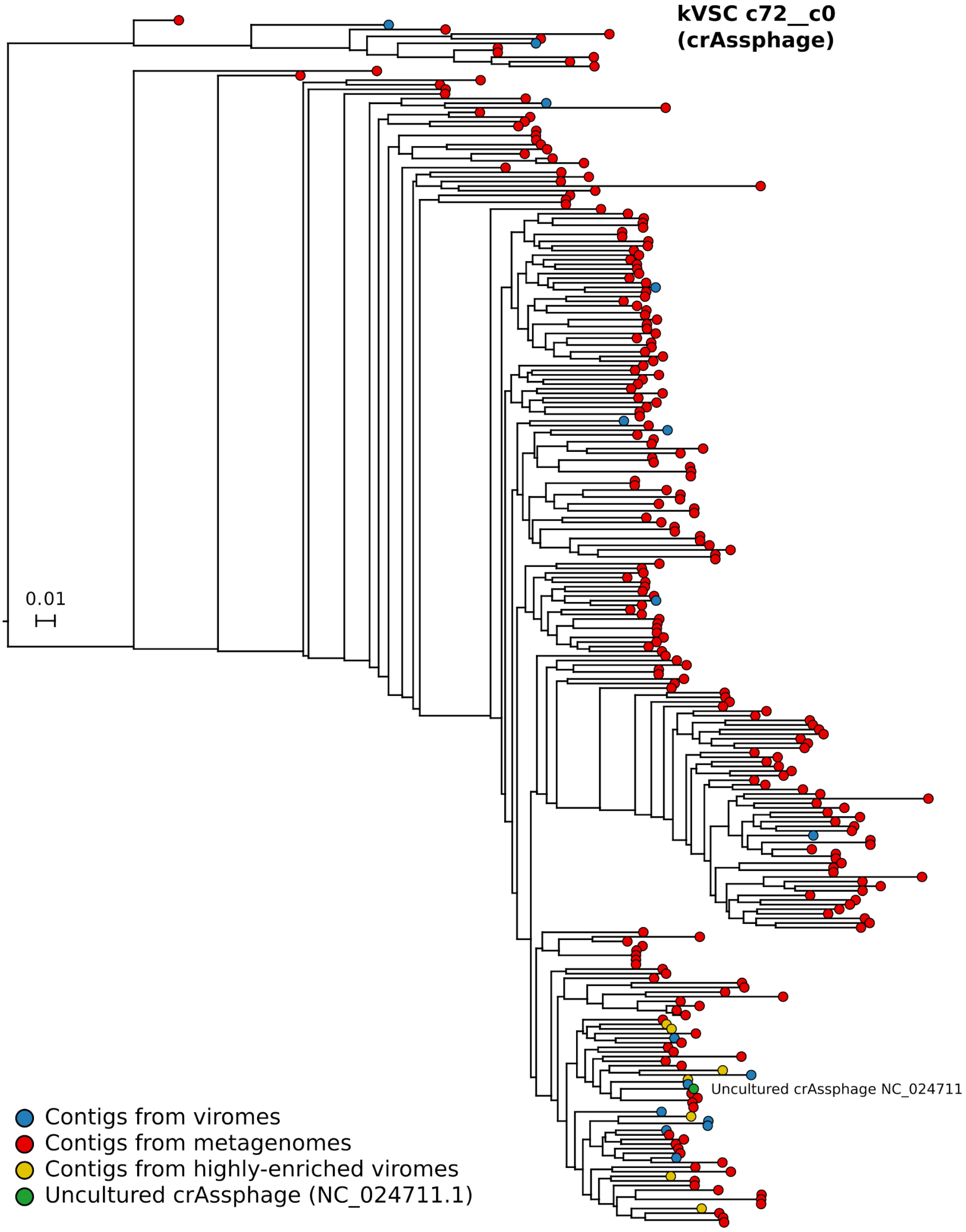


**Supplementary Figure 5. Phylogenetic tree of crAssPhage-like sequences (cluster c72__c0).** The 577 sequences in cluster c72__c0 were selected to build a phylogenetic tree. All sequences with a length outside 25% of the median length of all sequences in the cluster were excluded from the multiple-sequences alignment. The remaining 262 contigs were reconstructed from highly enriched viral contigs. Leaves are colored according to the origin of each sequence. Alignment statistics are provided in **Supplementary Table 6.**

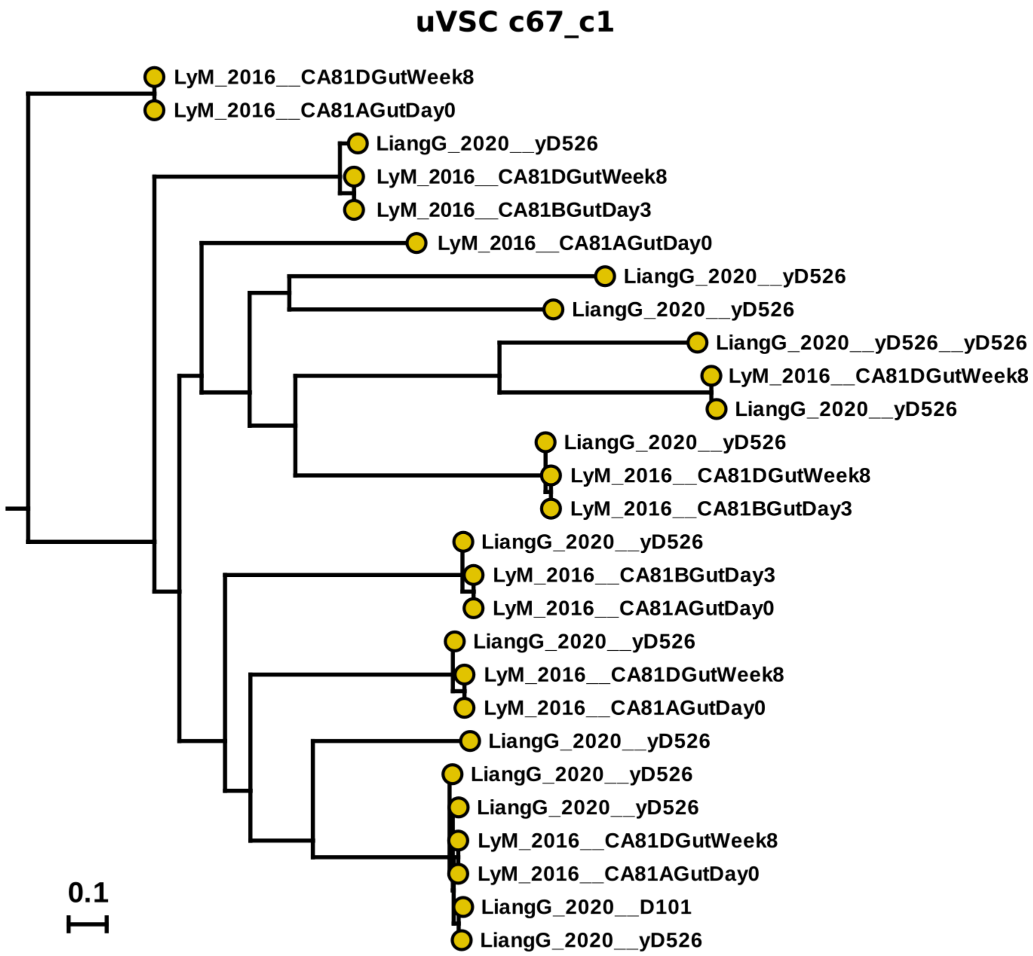


**Supplementary Figure 6. Phylogenetic tree of unknown viral sequence cluster c67_c1.** The 123 sequences in cluster c67__c1 were selected to build a phylogenetic tree of cluster c67__c1. Only sequences with a length within 15% of the median length of highly-enriched contigs in the cluster were kept for multiple-sequences alignment. All the 27 remaining sequences belonged to contigs reconstructed from highly enriched viral contigs. Median sequence length = 3,882 bp; Trimmed alignment length = 18,527 bp. Study ID and sample names are indicated in the label of each leaf. Alignment statistics are provided in **Supplementary Table 6.**

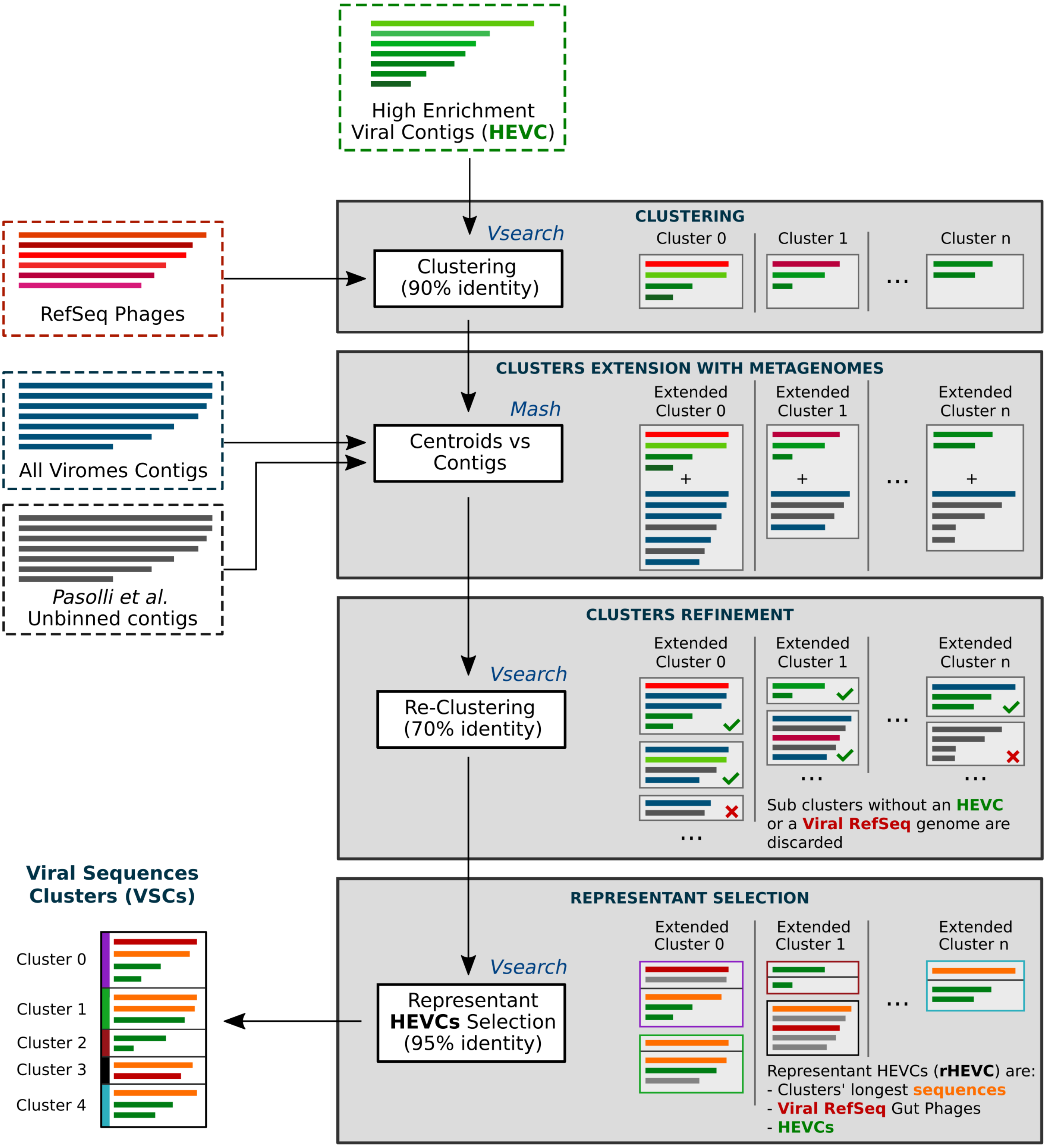


**Supplementary Figure 7 Detailed representation of the clustering procedure.** Contigs from highly enriched viromes (HEVCs, in green) were clustered together with gut bacteriophages reference genomes (in red). Clusters were extended by adding similar sequences from viromes (blue contigs) and unbinned metagenomes (gray contigs). After a second step of clustering at 70% identity, only clusters that contained at least one of the original sequences were kept. Finally, a third clustering step was performed to select the representative sequences of each cluster (Viral Sequences Clusters).

**Supplementary Table 1. Metadata and references of the 3,044 analyzed viromes.** The table contains metadata and references to the viromes from 49 datasets that were analyzed in this paper. References and sample counts aggregated by dataset are presented in **Tab. 1**. Detailed statistics for each virome are shown in **Tab. 2**.

**Supplementary Table 2. Annotation of the 120,041 contigs reconstructed from the metagenomic assembly of 255 highly enriched gut viromes.** Each contig was annotated against 80,853 bacterial reference genomes, as well as the MAGs from Pasolli *et al*. 2019, and the viral reference genomes in RefSeq.

**Supplementary Table 3. Selection of the 5,651 contigs from highly enriched viromes (HEVCs). Tab. 1** contains the full set of 5,651 contigs that were selected from gut viromes according to their prevalence and low contamination. Detailed information on the clusters in which each contig is available is reported in the first six columns. The results of viral-classification tools applied to each contig are also included. **Tab. 2** contains grouped statistics for each of the 3,944 clusters of sequences (see **Methods**).

**Supplementary Table 4. Prevalence of each Viral Sequence Cluster (VSC) in the surveyed metagenomes and viromes.** Prevalence of each Viral Sequence Cluster in the analyzed metagenomes (**Tab. 1**), viromes (**Tab. 2**), and all-samples (**Tab. 3**). Prevalence is defined as the percentage of samples in the dataset from which the cluster could be retrieved. Only samples where at least one VSC could be retrieved are included. The “percentage of datasets” column (4th and 10th columns of **Tabs. 1-3**), contains the percentage of datasets in which the VSC could be retrieved from at least one sample. **Tab. 4** contains the number of distinct samples in each dataset from which a VSC could be retrieved.

**Supplementary Table 5. CRISPR-spacers mapping statistics grouped by Viral Sequence Cluster (VSC). Tab. 1** shows the number of species associated with each VSCs (blast hits are considered if at < 3 SNPs distance, with min. Query coverage of 90% and min. Identity of 90%). **Tab. 2** shows the number of hits per species. **Tabs. 3-4** and **5-6** contain the same data grouped at genus and phylum level, respectively. **Tab. 7** contains the raw associations count.

**Supplementary Table 6. Multiple-Sequence Alignments Statistics.** Phylogenetic trees were built from multiple-sequence alignments filtered to maximize the phylogenetic signal (see **Methods**). The table recapitulates the number of sequences in each tree and the lengths of sequences before and after the alignment. The selection strategy indicates how the sequences of each cluster were selected. Sequences were kept if they had a length within 25% of the median of the lengths in the cluster (strategy “A”), or within the 15% of the median of the sequences from highly-enriched contigs (strategy “B”).

**Supplementary Table 7. Prevalence of the Viral Sequence Groups (VSGs) in 18,756 metagenomes.** A total of 18,756 human gut metagenomes from 81 datasets were mapped against the VSGs representatives to estimate the prevalence of each cluster in each dataset. Aggregated values are reported in **Tab. 1**. Per-dataset prevalences are reported in **Tab. 2.** Datasets names and references are reported in **Tabs. 3** and **4**.

**Supplementary Table 8. Phage-Host associations via CRISPR spacers.** Number of species and SGBs associated via CRISPR spacers to each of the 1,345 Viral Sequence Groups (VSGs, **Tab. 1**), and to each of the 45,872 representative sequences in the MetaPhlAn4 viral plugin (**Tab. 2)**.
